# SARS-CoV-2 nucleocapsid uniquely disrupts chromatin over pathophysiologically relevant gene promoters

**DOI:** 10.1101/2025.02.22.639619

**Authors:** Jane M Benoit, Jonathan H Dennis

**Affiliations:** Department of Biological Science, Florida State University, Tallahassee, FL, USA

## Abstract

SARS-CoV-2, the causative agent of COVID-19, is a positive-sense, single-stranded RNA virus that causes a spectrum of disease severity, from asymptomatic infection to severe illness to long-term sequelae. Similar to other human coronaviruses, SARS-CoV-2 proteins modulate host genomic responses through epigenomic modifications, facilitating viral replication and immune evasion. While the nucleocapsid protein is well known for its role in RNA stability and immune modulation, its impact on host chromatin organization remains unclear. To investigate this, we generated stable human alveolar cell lines expressing nucleocapsid proteins from endemic and pandemic human coronaviruses. Our analysis revealed that nucleocapsid proteins from all tested coronaviruses induced changes in nucleosome positioning and occupancy at specific gene promoters involved in coagulation pathways, hormone signaling, and innate immune responses. Additionally, SARS-CoV-2-specific alterations were identified in genes dysregulated in severe infections, suggesting a direct role for epigenomic modifications in disease pathophysiology. We also observed extensive changes in nucleosome susceptibility to nuclease digestion in SARS-CoV and SARS-CoV-2 samples that were not observed in common cold cell lines. Promoters with altered sensitivity and resistance to nuclease were linked to innate immune, metabolic, olfactory, and signaling pathways known to be dysregulated in severe COVID-19 and post-acute sequelae (PASC). These findings demonstrate that nucleocapsid protein expression alters chromatin structure at specific loci, implicating viral proteins in host genome dysregulation. Furthermore, we identified both shared and unique chromatin targets of SARS-CoV-2 and common cold coronaviruses, highlighting pathways for further investigation and potential therapeutic intervention.

**Importance:** Host chromatin is known to be modulated by coronaviruses during infections. However, the role of the nucleocapsid protein in these alterations are unknown. Here, we show that nucleocapsids from seven human coronaviruses alter nucleosome distribution and susceptibility to enzymatic digestion over specific gene promoters in a human lung cell line. Nucleocapsids from SARS-CoV and SARS-CoV-2 have the most prominent effects which are seen over genes involved in immune responses, metabolism, hormone signaling, and other pathways that are known to be dysregulated in severe COVID-19 and post-acute sequelae of COVID-19.

**Graphical abstract:** 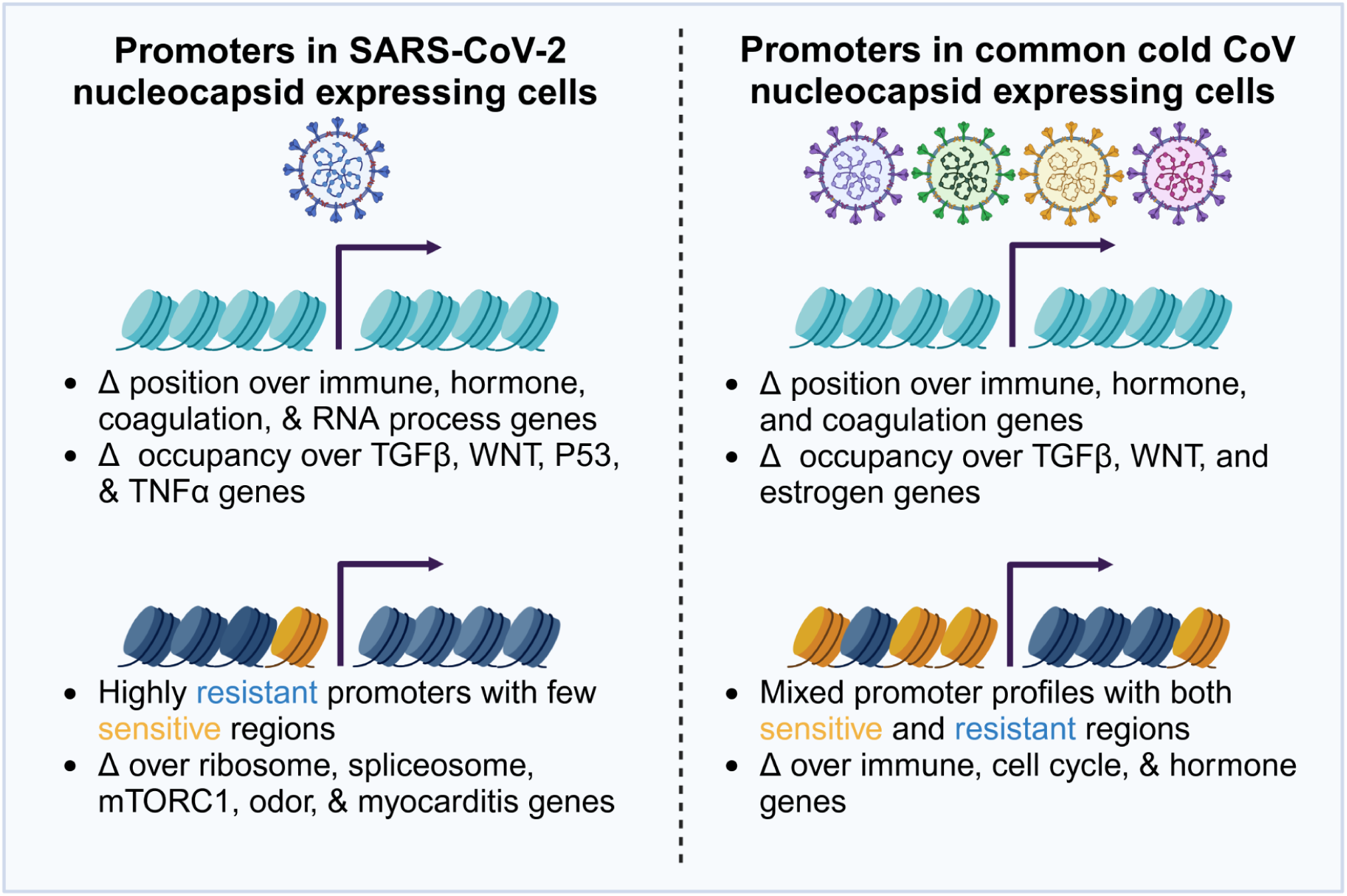

## Introduction

Coronaviruses are single-stranded positive-sense RNA viruses that infect a wide range of organisms including birds and mammals. Members of both the alphacoronavirus and betacoronavirus genera are able to infect humans and have been responsible for three human pandemics in the past two decades (1). There are four known human coronaviruses which are associated with common colds: HCoV-HKU1, HCoV-NL63, HCoV-OC43, and HCoV-229E (2–5). These viruses generally produce mild upper respiratory infections in healthy individuals, are collectively associated with approximately 30% of common colds (6, 7), and have a high seroprevalence in adults (8, 9). Both MERS-CoV and SARS-CoV can cause severe lower respiratory tract disease, leading to acute lung injury, respiratory failure, and death (10–16). In contrast, SARS-CoV-2, the causative agent of COVID-19 (17), produces a range of disease phenotypes from asymptomatic infections to severe disease with some individuals experiencing persistent symptoms in the months and years following infection (18, 19). Severe disease from SARS-COV-2 infection is mediated by dysregulation of immune responses (19–22) and the majority of the global population has been exposed to this virus. HCoV-NL63 and HCoV-229E are both alpha coronaviruses while HCoV-OC43, HCoV-HKU1, SARS-CoV, MERS-CoV, and SARS-CoV-2 are beta coronaviruses (23, 24). Each human coronavirus evolved in animal hosts and independently emerged as a human pathogen (25–27). A large reservoir of coronaviruses in animal populations presents a continued risk of zoonotic spillover into humans (10, 28–30).

Human coronavirus genomes are composed of a linear positive-sense single strand of RNA with two partially overlapping open reading frames encoding structural, accessory, and nonstructural proteins (31–33). The assembled viral particle contains an RNA genome which is coated with nucleocapsid protein and encased in an lipid bilayer embedded with spike, envelope, and membrane proteins (34, 35). Individual CoV proteins have specific functions in the cell which range from dampening host immune responses to promoting viral replication. CoV infection is known to directly interfere with host cells through disruption of host chromatin organization which leads to dysregulation of gene expression and can have long term consequences for the host cell and organism as a whole (36). Because chromatin architecture tightly regulates gene expression, viral disruption of these structures can profoundly impact cellular function

In eukaryotic cells, linear DNA is packaged within the nucleus as chromatin by wrapping DNA around a core octamer of eight histone proteins to form a nucleosome. Nucleosomes are joined together by linker DNA and are further compacted into chromatin fibers. This compaction compacts the DNA within the nucleus and regulates accessibility to repair, replication, and transcriptional machinery (37–40). Nucleosomes within promoters are especially critical for transcriptional regulation as they act as barriers transcription by preventing assembly of the pre-initiation complex (41) and depletion of nucleosomes results in aberrant expression of otherwise silent genes (42). Highly expressed genes are defined by highly phased nucleosomes flanking a nucleosome depleted region over the immediate transcription start site (TSS) (43–45). Epigenomic modifications of the underlying DNA and chromatin proteins contribute to this regulation without altering the underlying DNA sequence. These modifications include DNA methylation (46–48), histone modifications such as acetylation, methylation, phosphorylation or ubiquitination (49–51), and RNA modifications (52–54). Each of these modifications interact with the overall chromatin architecture and contribute to regulation of gene expression.

Coronaviruses have evolved mechanisms to perturb host chromatin to promote viral replication and limit immune responses on both local and genome wide scales. SARS-CoV and MERS-CoV infection result in altered or delayed interferon gene responses which are regulated through alterations in histone post translational modifications over specific immune response genes (36, 55). HCoV-229E infection also results in altered histone modifications and pol II recruitment which are associated with differential gene expression (56). Alterations in histone post translational modifications extend to transposable elements which are uniquely modulated following SARS-CoV-2 infection (11). Recent high-throughput CRISPR screens have identified chromatin remodelers, cohesin, and transcription factors as key regulators of host responses to infection, underscoring the role of chromatin organization in viral pathogenesis (57–59). Chromatin accessibility, as measured by ATACseq, is disrupted over immune genes in peripheral blood mononuclear cells from recovering COVID-19 patients (60) and those with severe and mild disease (61, 62) indicating chromatin alterations occur in a variety of disease contexts. SARS-CoV-2 infection alters higher-order chromatin structure, leading to restructuring of HiC A/B compartments. This is accompanied by reduced loop extrusion and increased local variability over immune response genes. These changes are coupled with alterations in HPTM deposition and gene expression (63, 64). Interestingly, these higher order modifications were not observed in infection with HCoV-OC43 (64) but similar patterns of compartment mixing and loss of stability over pro-inflammatory genes are observed in HCoV-229E infection (65) suggesting variability across CoVs. Specific large scale chromatin rearrangements in olfactory cells have been associated with anosmia following SARS-CoV-2 infection in hamsters and humans (66) suggesting functional implications for these chromatin alterations. Individual SARS-CoV-2 proteins have known impacts on host chromatin structure and function. The ORF8 protein functions as a histone mimic (67) and disrupts host gene regulation disrupting histone post translational modifications. NSP1 is a multifunctional protein which disrupts host mRNA translation and stability (68–70), blocks nuclear export (71, 72), antagonizes type I interferon signaling (73), and reduces Pol II occupancy and silences immune genes (74, 75). Other viral proteins have known functions on RNA rather than chromatin such as SARS-CoV-2 nucleocapsid which interacts with genes involved in RNA synthesis, splicing, and stability (76). While prior studies have examined the impact of CoV infections on host chromatin, the direct effects of the majority of individual viral proteins on nucleosome architecture remain underexplored.

Despite the global importance of coronaviruses and the ongoing risk of future spillovers, their direct effects on chromatin architecture at the nucleosomal scale—one of the most fundamental levels of gene regulation—remain poorly understood. To answer this question, we generated a reductionist system to study the impacts of seven human CoV nucleocapsid proteins on nucleosome architecture in a human lung cell line. We opted to study the nucleocapsid protein as it is highly abundant, generally conserved, and has known roles in RNA regulation but is understudied in the context of chromatin (77). Nucleocapsid contributes to viral RNA replication and stability (78–81) and has broad RNA binding capabilities (82–84). The SARS-CoV-2 nucleocapsid protein induces expression of *Il-6* and other inflammatory cytokines through *NF-Kb* signaling (85, 86), inhibits prevents RIG-I recognition of viral dsRNA (87, 88), and prevents IRF3 phosphorylation to limit type I interferon responses (89). Our reductionist system allowed us to probe the effects of nucleocapsid on chromatin structure which might be overshadowed in an infection model due to wide ranging effects on host cells.

To determine the role of CoV nucleocapsid proteins on nucleosome distribution and susceptibility to enzymatic digestion within promoters, we generated stable A549 human lung epithelial cell lines expressing the full length nucleocapsid proteins of seven human CoVs. Replicate cell lines were generated for each of the following nucleocapsids: HCoV-HKU1, HCoV-NL63, HCoVOC43, HCoV-229E, SARS-CoV, MERS-CoV, and SARS-CoV-2 which were compared to cells expressing an empty vector construct. The distribution of nucleosomes across promoters was determined by digesting crosslinked chromatin with Micrococcal nuclease (MNase), an enzyme which selectively cleaves linker DNA leaving nucleosomal DNA intact for high throughput sequencing (43, 90). In addition to mapping the location of individual nucleosomes, a titration of MNase was used to determine the susceptibility of a given nucleosome to enzymatic digestion (91–96). This method identifies nucleosomes which are sensitive or resistant to enzymatic digestion and is correlated with gene expression and transcription factor binding suggesting that it is a useful method for identification of active and silent genes. We chose to focus our analysis on promoter regions as nucleosomes within promoter regions have distinct roles in gene regulation, are highly occupied, and are especially well positioned compared to the rest of the genome.

Our results indicate that SARS-CoV-2 nucleocapsid expression specifically alters nucleosome positioning and occupancy at genes involved in immune response, coagulation, and hormone signaling—pathways linked to severe COVID-19 outcomes and post-acute sequelae. Many of these genes are differentially expressed in severe acute disease and continue to be dysregulated in post acute sequelae (PASC) months following infection. In contrast, we identified greater disruptions in susceptibility to nuclease digestion with unique alterations occurring in both common cold and pandemic viruses. SARS-CoV-2 specific alterations were localized over genes associated with tight junction interactions, innate immune signaling, and endocrine pathways. Our findings reveal distinct alterations in nucleosome distribution and nuclease sensitivity upon expression of SARS-CoV-2 nucleocapsid in a controlled reductionist system. Affected genes were associated with disease phenotypes and may offer insight into the pathophysiology of both common cold and pandemic CoVs.

## Results

### Nucleosome distribution over promoters is largely stable in cells expressing CoV nucleocapsids

To investigate the impact of coronavirus (CoV) nucleocapsid proteins on nucleosome architecture, we generated A549 human lung alveolar cell lines expressing nucleocapsid genes from seven human CoVs: 229E, NL63, HKU1, OC43, MERS-CoV, SARS-CoV, and SARS-CoV-2. While these nucleocapsid proteins share conserved functions, they exhibit distinct amino acid sequences (Supp. Fig. 1) and can be classified into alphacoronaviruses, common cold betacoronaviruses, and pandemic betacoronaviruses (Fig. 1A). To ensure the correct integration of full-length wild-type nucleocapsid genes, we validated all cell lines through Sanger sequencing.

**Figure 1.**
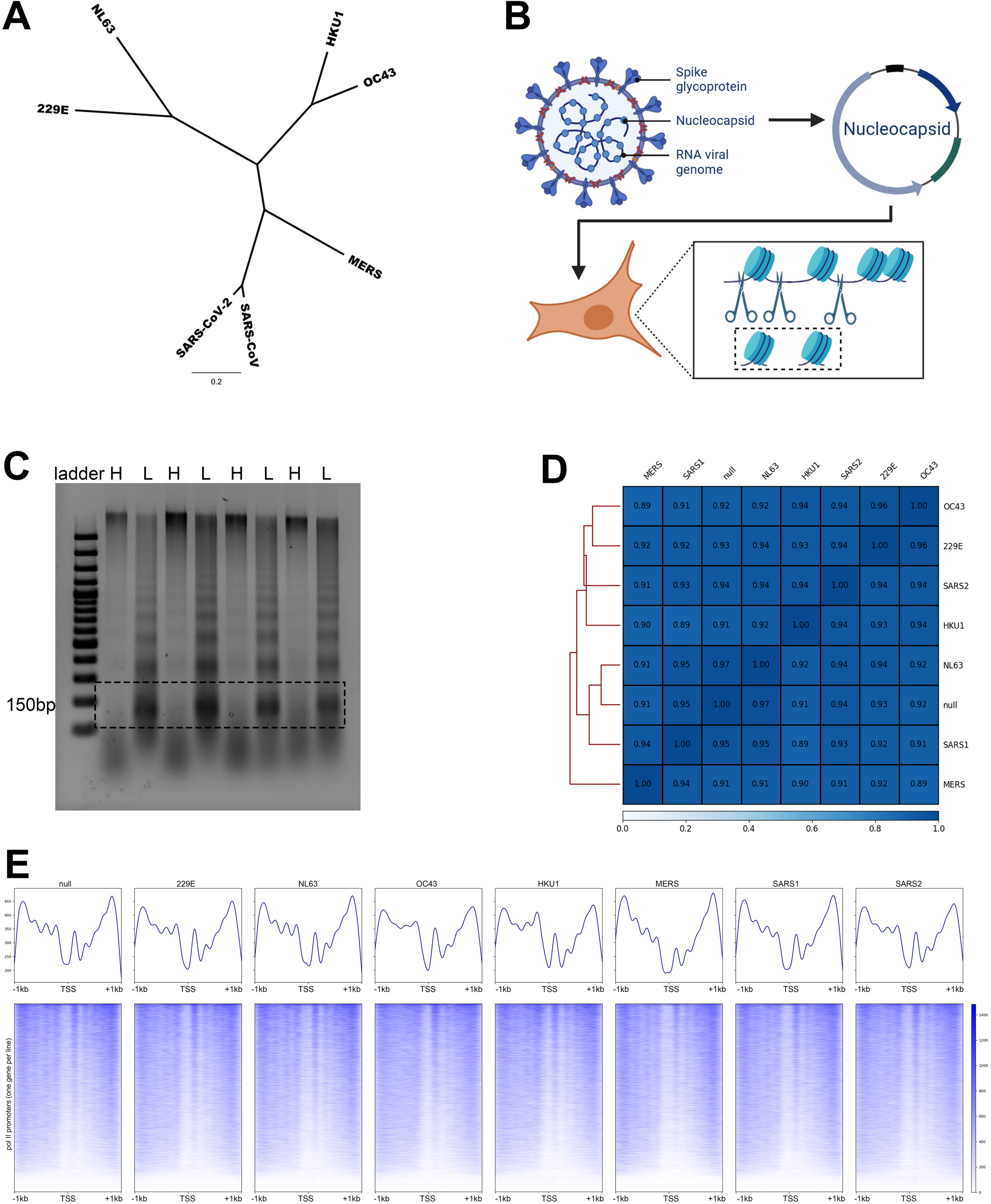
Nucleosome distribution over promoters in CoV nucleocapsid expressing cell lines is highly similar. A. Phylogenetic tree of CoV nucleocapsid proteins generated with a Jukes-Cantor genetic distance model and neighbor joining. Common cold alphaCoVs (229E & NL63), common cold betaCoVs (HKU1 & OC43), and pandemic betaCoVs (MERS, SARS-CoV, & SARS-CoV-2) cluster together. B. Schematic of CoV structure showing nucleocapsid protein. Nucleocapsid genes from six CoVs were engineered into plasmid vectors and expressed in A549 cells. Chromatin from stable cell lines was digested with MNase and mononucleosomal DNA was used to prepare sequencing libraries for mapping experiments. C. Representative agarose gel showing nucleosomal ladders following light (L) ad heavy (H) digests with MNase. D. Spearman correlation between nucleosome signal across 2kb regions of all pol II promoters divided into 100bp bins. Correlation coefficients are listed within each square of the heatmap. E. Heatmaps of nucleosome distribution (blue) over a 2kb region centered on the TSS of pol II promoters (one gene per line) with a line plot showing average signal above each plot. The overall gene order is identical across heatmaps and is determined by maximum signal values.

To assess nucleosome positioning, we digested crosslinked chromatin with Micrococcal nuclease (MNase) and isolated mononucleosome-sized DNA fragments for nucleosome mapping (workflow shown in Fig. 1B). A combination of light and heavy MNase digests allowed us to capture nucleosomes with varying susceptibility to nuclease cleavage (Fig. 1C). Using a previously validated promoter sequence capture approach (97, 98), we enriched for nucleosomes within 2 kb of RefSeq transcription start sites (TSSs), achieving strong enrichment over Pol II promoters with robust coverage from libraries containing as few as 20 million reads per sample (Supp. Fig. 2).

Comparative analysis revealed that nucleosome distribution across Pol II promoters was highly similar between CoV nucleocapsid-expressing and empty vector control cell lines. The Spearman’s correlation coefficient exceeded 0.89 for nucleosome distribution signals across 100-bp bins, indicating high overall similarity between samples (Fig. 1D). Furthermore, aggregate nucleosome maps across all samples (Fig. 1E) demonstrated a conserved promoter architecture, including: 1. Four weakly phased but highly occupied upstream nucleosomes 2. A prominent nucleosome-depleted region (NDR) over the TSS 3. A strongly positioned and highly occupied +1 nucleosome 4. Four weakly phased but highly occupied nucleosomes downstream of the TSS. Although variability was observed in the phasing and occupancy of individual nucleosomes, the overall promoter nucleosome architecture remained consistent across conditions. These patterns align with previously reported human nucleosome distribution maps (43, 93, 94, 97, 98), supporting the robustness of our findings.

### Nucleosome positioning and occupancy is altered over specific gene promoters in cells expressing CoV nucleocapsids

Although overall nucleosome distribution remained highly similar across samples, statistical analysis revealed CoV-specific alterations in nucleosome positioning and occupancy when compared to empty vector controls. Nucleosome positioning refers to the location of a nucleosome relative to the underlying DNA, whereas occupancy is defined by the summit value of the nucleosome peak. To identify changes in positioning, we used DANPOS3 (102) to compare nucleosome positions in CoV nucleocapsid-expressing cell lines with those in empty vector controls. High-confidence repositioning events were defined as a shift of >80 bp in summit position which was consistent across replicates. Across conditions, we identified between 119 to 361 genes with at least one repositioned nucleosome in their promoter regions (Fig. 2A, Table 2). Notably, the majority of repositioned genes were unique to each CoV nucleocapsid, with limited overlap between common cold CoVs and pandemic CoVs (Fig. 2A, Supp. Table 1).

**Figure 2.**
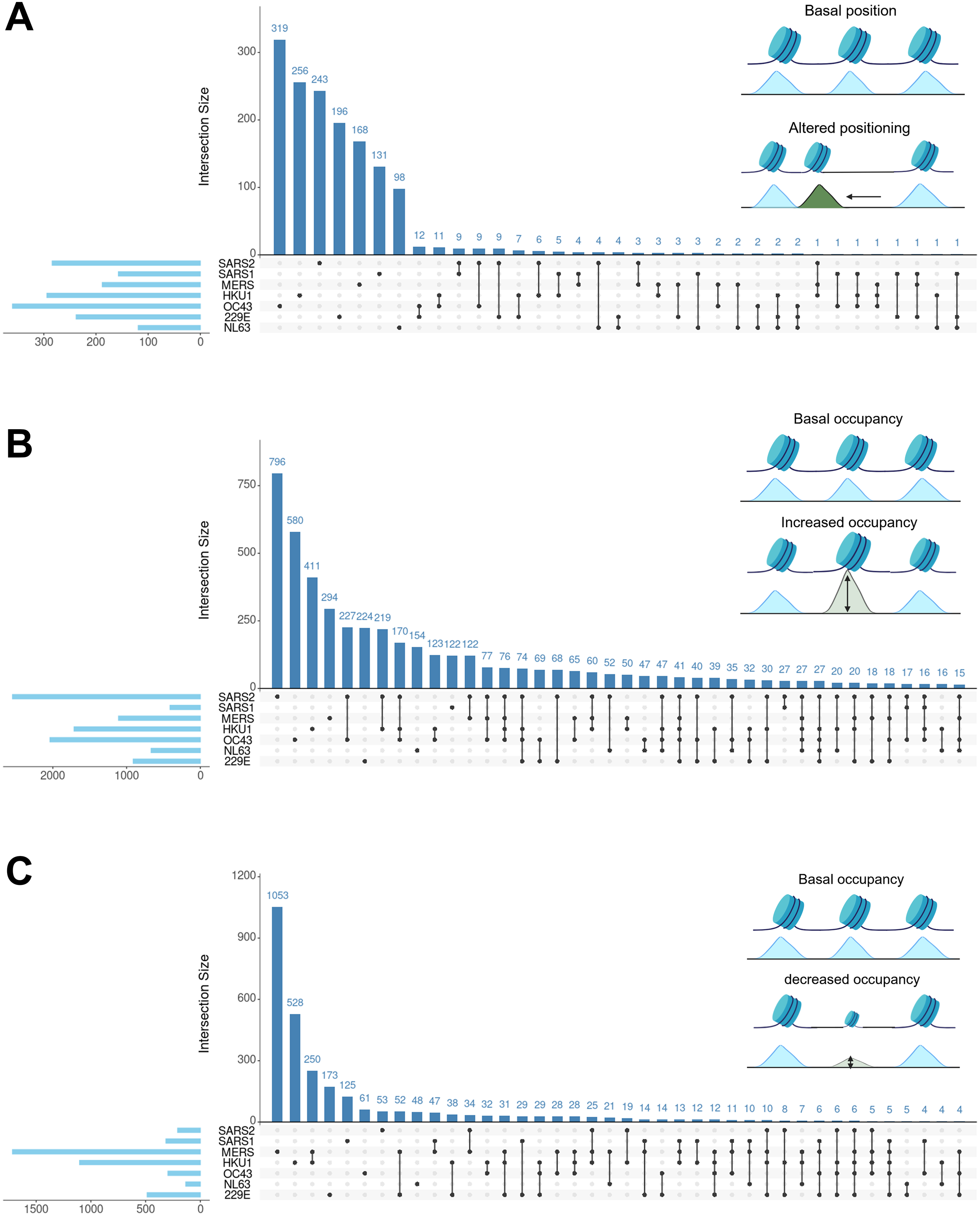
Selected promoters in CoV nucleocapsid expressing cells have redistributed nucleosomes. A. Upset plot showing the number of shared and unique genes with one or more nucleosomes with alterations in position (greater than 80bp) across samples. The pale blue horizontal bars show the total number of repositioned nucleosomes while vertical bars and connecting dots show shared genes across samples. B. The same for genes with increased occupancy signal in one or more nucleosomes per gene. C. The same for genes with decreased occupancy signal in one or more nucleosomes per gene.

**Table 1.** Plasmid constructs used to generate stable cell lines.

| Addgene # | Plasmid name | insert | Antibiotic resistance |
| --- | --- | --- | --- |
| 151912 | pGBW-m4134902 | 229E nucleocapsid | puromycin |
| 151960 | pGBW-m4134906 | OC43 nucleocapsid | puromycin |
| 151957 | pGBW-m4134908 | NL63 nucleocapsid | puromycin |
| 151948 | pGBW-m4134897 | HKU1 nucleocapsid | puromycin |
| 151951 | pGBW-m4134903 | SARS2-CoV-2 nucleocapsid | puromycin |
| NA | pTwist_SARS1 | SARS-CoV-1 nucleocapsid | puromycin |
| NA | pTwist_MERS | MERS-CoV nucleocapsid | puromycin |
| NA | pTwist_null | Empty vector | puromycin |

**Table 2.** Number of genes with one or more re-distributed nucleosomes within 2kb promoter regions in each CoV nucleocapsid cell line compared to empty vector (null).

| Time | repositioning | increased occupancy | decreased occupancy |
| --- | --- | --- | --- |
| 229E | 238 | 910 | 486 |
| NL63 | 118 | 667 | 133 |
| HKU1 | 294 | 1712 | 1103 |
| OC43 | 360 | 2039 | 296 |
| MERS | 188 | 1109 | 1716 |
| SARS1 | 157 | 411 | 314 |
| SARS2 | 284 | 2547 | 207 |

Among genes with nucleosome repositioning unique to SARS-CoV, we identified PABPC1, a gene involved in RNA transport, translation, and stability. The Polyadenylate-binding protein 1 (PABPC1) has been previously reported to interact with the SARS-CoV-2 nucleocapsid (76, 109). Gene ontology (GO) analysis of promoters with repositioned nucleosomes across CoV nucleocapsid conditions revealed associations with innate immune signaling, coagulation, hormone responses, and signaling pathways (Supp. Fig. 3). Additionally, many of these genes overlap with known dysregulated genes in SARS-CoV-2 infection and severe disease progression, further supporting a potential functional relationship between CoV nucleocapsid expression, chromatin remodeling, and pathophysiological outcomes.

We next analyzed nucleosome occupancy changes, defining significant alterations as a ≥5-fold increase or decrease compared to empty vector controls, with an FDR and p-value <0.05. High-confidence altered occupancy events were filtered for consistency across replicates. This analysis identified between 412 to 2,548 genes with increased nucleosome occupancy (Fig. 2B, Table 2) and between 134 to 1,717 genes with decreased nucleosome occupancy compared to controls (Fig. 2C, Table 2). Similar to repositioning, most nucleosome occupancy changes were unique to a single nucleocapsid condition, with moderate overlap between common cold and pandemic CoVs (Fig. 2, Supp. Table 2). GO analysis of promoters with increased nucleosome occupancy revealed enrichment in pathways related to TGF-β signaling, WNT signaling, Hedgehog signaling, and estrogen responses (Supp. Fig. 4). These pathways align with previous studies linking these pathways to lung fibrosis, sex differences in disease severity, and other disease pathologies (110–113). Promoters with increased occupancy in SARS-CoV-2-expressing cells also included PABPC1 and LARP1, both of which have established interactions with SARS-CoV-2 nucleocapsid protein, as well as 36 additional genes known to interact with other SARS-CoV-2 proteins (76, 109).

For genes with decreased nucleosome occupancy, GO analysis identified many of the same pathways as those with increased occupancy, along with p53 signaling and TNF-α pathways (Supp. Fig. 5, Supp. Table 3). TNF-α signaling is a major driver of cytokine storm and severe COVID-19 outcomes, reinforcing a potential link between CoV nucleocapsid expression, chromatin remodeling, and immune dysregulation (21, 113–117). Collectively, these results suggest that CoV nucleocapsid proteins induce targeted nucleosome distribution changes at specific gene promoters, many of which are functionally relevant to host response pathways, disease progression, and severe COVID-19 outcomes.

**Table 3.** Number and percent of sensitive and resistant nucleosomes within promoters in each CoV nucleocapsid cell line. Nucleosomes with a log2 fold change in summit score greater than 1.5 combined with a p-value and FDR under 0.05 were classified as sensitive if more abundant in light digests and resistant if more abundant in heavy digests. All nucleosomes not meeting these thresholds were classified as neutral.

| time | # sensitive | # resistant | # neutral | % sensitive | % resistant |
| --- | --- | --- | --- | --- | --- |
| null | 25767 | 19856 | 131928 | 14.51 | 11.18 |
| 229E | 28386 | 30888 | 114402 | 16.34 | 17.78 |
| NL63 | 38165 | 34514 | 84282 | 24.31 | 21.99 |
| HKU1 | 49148 | 47324 | 68139 | 29.86 | 28.75 |
| OC43 | 29116 | 28543 | 110073 | 17.36 | 17.02 |
| MERS | 16562 | 15736 | 125166 | 10.52 | 9.99 |
| SARS1 | 30916 | 20715 | 114131 | 18.65 | 12.50 |
| SARS2 | 41455 | 49218 | 66946 | 26.30 | 31.23 |

### Nucleosome susceptibility to nuclease digestion is associated with active promoters

While nucleosome positioning and occupancy provide key insights into chromatin alterations following CoV nucleocapsid expression, these analyses alone do not fully capture the breadth of pathways affected by CoV infection. Because nucleosome distribution mapping offers a largely static view of chromatin organization, we expanded our analysis to include nucleosome susceptibility to nuclease digestion to better understand gene regulation in infected cells. Using a titration of Micrococcal Nuclease (MNase), we generated light and heavy digests to identify nucleosomes with differential susceptibility to nuclease activity (Fig. 1C). Nucleosomes preferentially released under light digest conditions were classified as “sensitive,” whereas those requiring heavy digestion were labeled “resistant” (summarized in Fig. 3A). This approach has been widely used to characterize promoter chromatin landscapes and distinguish between active and silent genes across various organisms and conditions (91–96).

**Figure 3.**
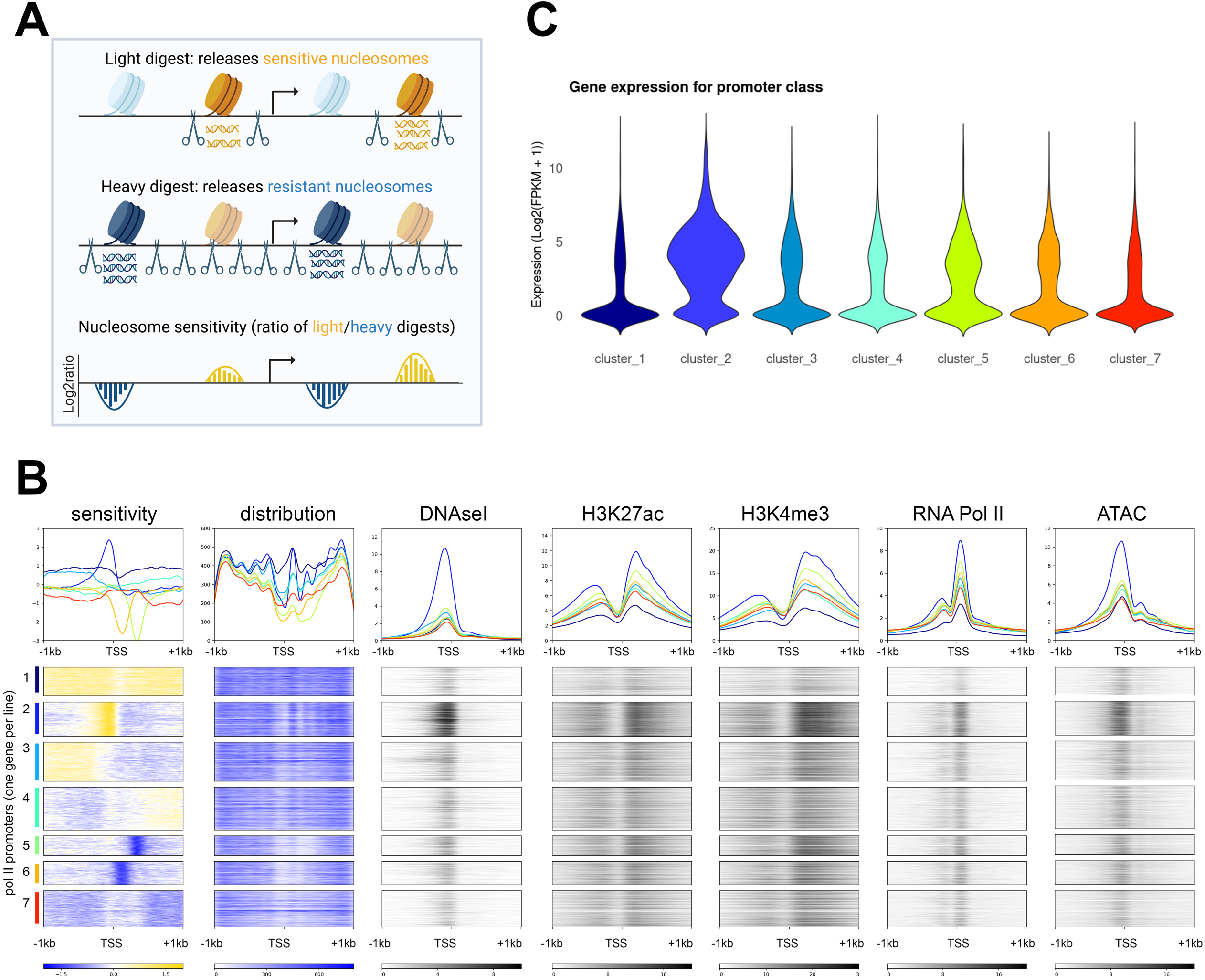
Nucleosome susceptibility to nuclease digestion is associated with active chromatin marks and increased gene expression. A. Schematic of MNase sensitivity assay. A subset of sensitive (gold) nucleosomes are preferentially released under light MNase digest conditions while a different population of resistant (blue) nucleosomes are released under heavy MNase digest conditions. The log2ratio of light over heavy nucleosome signal results in positive values for sensitive nucleosomes and negative values for resistant nucleosomes. B. Heatmap of nucleosome sensitivity (gold) and resistance (blue) over 2kb regions of pol II promoters centered on the TSS for the empty vector sample. Dominant promoter classes were identified by kmeans clustering and gene order is applied to all subsequent heatmaps. Heatmaps of nucleosome distribution signal (blue), DNAseI hypersensitivity signal (black), H3K27ac signal (black), H3K4me3 signal (black), RNA Pol II signal (black), and ATAC signal (black) are sorted by the same gene order as the sensitivity map. Line plots showing the average signal are shown above each heatmap. C. Violin plot of gene expression (log2 (FPKM +1)) for all genes within each sensitivity cluster defined in B.

To establish a baseline chromatin landscape, we analyzed empty vector control cells and assessed the relationship between nucleosome susceptibility and key epigenomic features of active chromatin. Using publicly available ENCODE datasets (107, 108), we compared nucleosome sensitivity patterns with DNase I hypersensitivity, histone post-translational modifications (PTMs), RNA polymerase II (Pol II) occupancy, and ATAC-seq accessibility (Fig. 3B). This analysis identified several distinct classes of promoters based on their nucleosome sensitivity patterns. Promoters in cluster 1 exhibited diffuse sensitivity, characterized by weakly phased nucleosomes, an occupied nucleosome-depleted region, and reduced levels of H3K27ac, H3K4me3, Pol II, and ATAC-seq signals (Fig. 3B). These promoters were poorly expressed (Fig. 3C), consistent with their repressed chromatin state.

In contrast, cluster 2 promoters displayed a strongly positioned sensitive region over the transcription start site (TSS), a well-defined nucleosome-depleted region, and a strongly positioned +1 nucleosome. This class was hypersensitive to DNase I and exhibited significantly higher levels of H3K4me3, H3K27ac, Pol II, and ATAC-seq signal compared to other clusters (Fig. 3B). As expected given the convergence of active chromatin marks, these promoters were highly expressed (Fig. 3C). These findings suggest that nucleosome sensitivity can be broadly categorized into two functional patterns: diffuse sensitivity, associated with low transcriptional activity, and localized sensitivity over the TSS, linked to highly active promoters.

Both mixed (clusters 3 and 4) and resistant (clusters 5-7) promoter groups showed defined nucleosome-depleted regions and moderate levels of active histone PTMs, Pol II, and ATAC-seq signals (Fig. 3B). These clusters exhibited similar expression levels, with slightly higher transcriptional activity observed in positioned resistant promoters compared to mixed clusters (Fig. 3C). Together, these findings demonstrate that nucleosome susceptibility to MNase digestion distinguishes between active and silent chromatin states. This analysis suggests that nucleosome susceptibility to nuclease digestion can distinguish between active and silent chromatin states. These findings provide a framework for interpreting how alterations in nucleosome sensitivity following CoV nucleocapsid expression may contribute to transcriptional regulation and disease progression.

### The majority of promoters in CoV nucleocapsid expressing cells exhibit altered susceptibility to nuclease digestion

To assess the impact of CoV nucleocapsid expression on chromatin accessibility, we examined nucleosome susceptibility to nuclease digestion across all nucleocapsid-expressing cell lines. Unlike nucleosome distribution, which remained relatively consistent, susceptibility to nuclease digestion was highly variable across cell lines. Pairwise Spearman’s correlations over promoter regions ranged from 0.24 to 0.57, indicating substantial differences in chromatin accessibility between conditions (Fig. 4A). When mapping susceptibility across all Pol II promoters, we observed widespread disruption of both sensitive and resistant regions (Fig. 4B). In empty vector control cells, promoter susceptibility followed distinct patterns, including diffusely sensitive, highly positioned sensitive, mixed, positioned resistant, and diffusely resistant groups (clusters 1–7). These patterns were largely preserved in common cold CoVs but were severely disrupted in MERS, SARS-CoV, and SARS-CoV-2 nucleocapsid-expressing cells (Fig. 4B).

**Figure 4.**
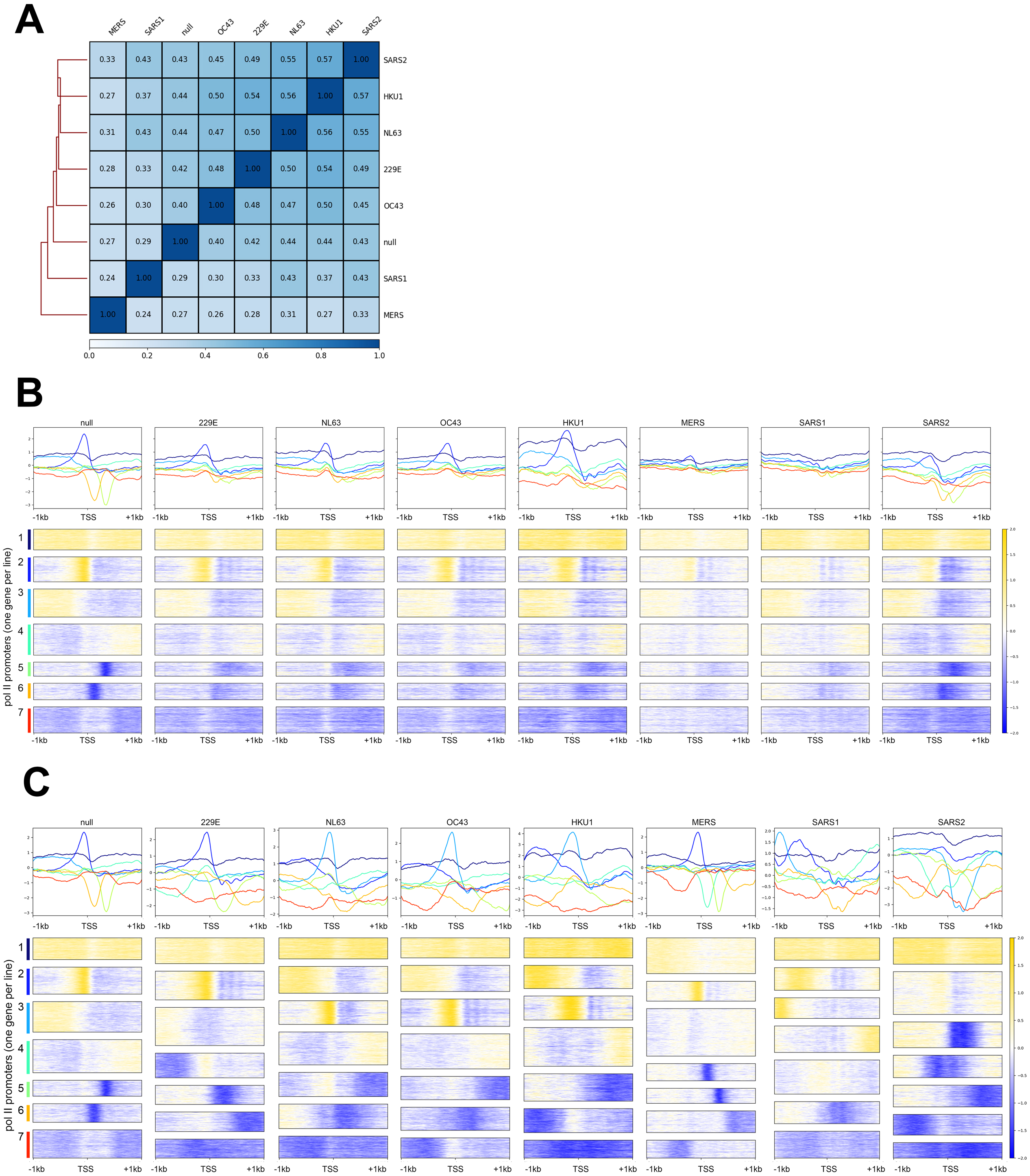
The majority of promoters in CoV nucleocapsid expressing cells have altered susceptibility to nuclease digestion. A. Spearman correlation of nucleosome sensitivity signal over 2kb regions of all pol II promoters divided into 100bp bins. Correlation coefficients are listed within each square of the heatmap. B. Heatmap of nucleosome sensitivity (gold) and resistance (blue) over 2kb regions of pol II promoters centered on the TSS for all samples. Line plots showing the average signal are shown above each heatmap. Kmeans clustering of the empty vector sample was used to establish gene sort order for all samples. C. Heatmap of nucleosome sensitivity (gold) and resistance (blue) over 2kb regions of pol II promoters centered on the TSS for all samples. Kmeans clustering was used to independently generate gene groups for each individual sample. Gene order is accordingly different in each heatmap.

This disruption did not result from a global loss of susceptibility but rather from nucleosomes transitioning between sensitive, neutral, and resistant states at specific genomic locations. Quantification of the proportion of sensitive and resistant nucleosomes in each sample confirmed that nucleocapsid-expressing cell lines retained similar or even greater levels of sensitivity and resistance compared to the empty vector control (Table 3). Independent clustering of each sample revealed the emergence of new susceptibility patterns in MERS, SARS-CoV, and SARS-CoV-2 conditions (Fig. 4C). While common cold CoV samples largely maintained the prominent promoter classes seen in control cells, MERS-expressing cells displayed an expanded diffusely sensitive promoter group and a reduction in mixed-susceptibility promoters (Fig. 4C, compare MERS to null). SARS-CoV-expressing cells exhibited a loss of resistant nucleosomes, an increase in mixed promoter classifications, and the disappearance of the highly positioned sensitive cluster over the TSS (Fig. 4C, compare SARS-CoV to null). Similarly, SARS-CoV-2-expressing cells lacked the TSS-sensitive cluster and showed an increased proportion of strongly resistant promoters (Fig. 4C, compare SARS-CoV-2 to null). Together, these findings indicate that CoV nucleocapsid expression alters nucleosome susceptibility in a virus-specific manner, with the most pronounced disruptions occurring in MERS, SARS-CoV, and SARS-CoV-2 conditions.

To further investigate the functional consequences of these chromatin alterations, we tracked the movement of genes between promoter classes across conditions. The majority of genes within the diffusely sensitive and silent promoter class were unique to the MERS-expressing cells, reflecting the expansion of this cluster (Fig. 5A, compare MERS cluster 1, Supp. Table 4). The second-largest gene group was common across all conditions, with gene ontology (GO) analysis revealing enrichment for pathways related to coagulation cascades and interferon-gamma responses (adj. p-value < 0.05, MSigDB Hallmark GO database). Additionally, 282 genes in this promoter class were unique to SARS-CoV-2 and were enriched for GO terms associated with olfactory transduction and viral myocarditis (KEGG GO database). Notably, genes transitioning from active mixed or resistant promoter classes in control cells to silent promoter classes in SARS-CoV-2 samples suggest potential transcriptional downregulation driven by SARS-CoV-2 nucleocapsid expression.

**Figure 5.**
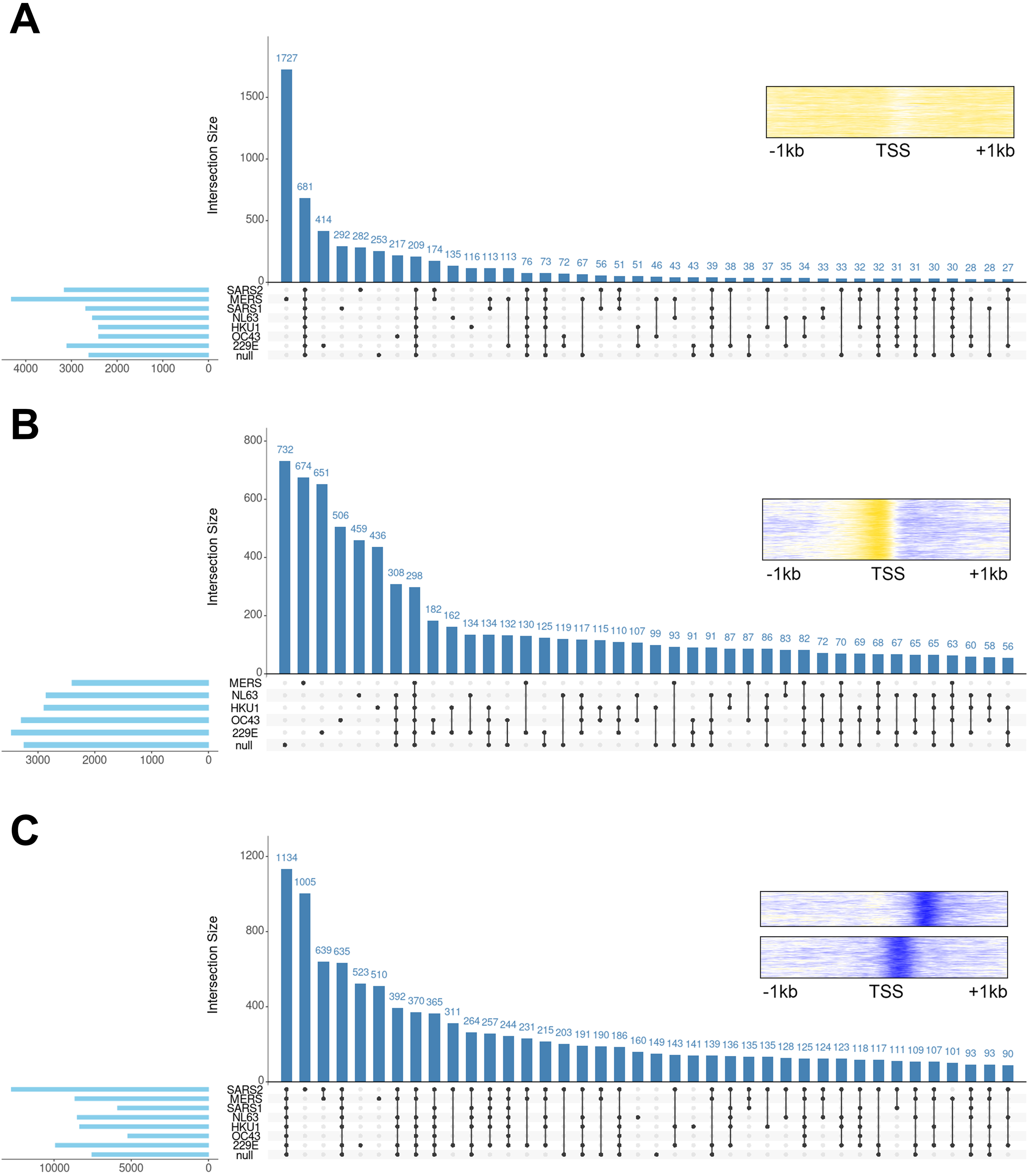
Nucleosome susceptibility to MNase is variable across CoV nucleocapsid expressing cell lines. A. Upset plot showing the number of shared and unique genes classified as having diffusely sensitive promoters. The pale blue horizontal bars show the total number of genes in each sample while vertical bars and connecting dots show shared genes across samples. B. Upset plot showing the number of shared and unique genes in the promoter class dominated by a highly positioned sensitive region over the immediate TSS. The pale blue horizontal bars show the total number of genes in each sample while vertical bars and connecting dots show shared genes across samples. C. Upset plot showing the number of shared and unique genes in resistant promoter classes. The pale blue horizontal bars show the total number of genes in each sample while vertical bars and connecting dots show shared genes across samples.

The highly expressed TSS-sensitive promoter class was absent in SARS-CoV and SARS-CoV-2 samples. Genes within this class were largely condition-specific, but a subset was shared across all common cold CoVs (Fig. 5B, Supp. Table 5). GO analysis showed significant enrichment for ribosome- and spliceosome-related functions, as well as coronavirus disease pathways (adj. p-value < 0.005, KEGG GO database). The absence of this cluster in SARS-CoV and SARS-CoV-2 samples suggests a distinct chromatin regulatory mechanism over these promoters, potentially contributing to gene repression in these conditions.

For subsequent analyses, all resistant promoter classes were combined due to their similar gene expression and epigenomic profiles. The majority of genes within resistant promoter classes were shared across all conditions (Fig. 5C, Supp. Table 6) and were significantly enriched for pathways associated with apical junctions, KRAS signaling, and myogenesis (MSigDB Hallmark GO database). However, SARS-CoV-2-expressing cells exhibited an expansion of resistant promoter classes, leading to an increase in SARS-CoV-2-specific genes within this group. These genes were enriched for GO terms related to Myc targets, protein secretion, oxidative phosphorylation, and mTORC1 signaling (MSigDB Hallmark GO database). Overall, these findings indicate that CoV nucleocapsid expression induces widespread alterations in nucleosome susceptibility to nuclease digestion, with potential consequences for transcriptional regulation and disease phenotypes. The pronounced disruptions observed in MERS, SARS-CoV, and SARS-CoV-2 samples suggest virus-specific mechanisms of chromatin remodeling, which may underlie differential pathogenicity and host responses.

## Discussion

Human coronavirus (CoV) infections are known to induce epigenomic alterations in host cells, including modifications to histone post-translational modifications (11, 36, 55, 56), altering accessibility as measured by ATACseq (60–62), and disrupting higher order chromatin architecture (63–66). These alterations influence gene expression and may contribute to both short- and long-term cellular dysfunction, even after viral clearance. Individual viral proteins can affect chromatin biology by acting as histone mimics (67), disrupting signaling pathways leading to transcription factor translocation to the nucleus (73, 88, 89, 118), altering RNA stability (68–70, 76, 78), and altering pol II occupancy (74, 75). Here, we systematically compare the effects of human CoV nucleocapsids on nucleosome distribution and susceptibility to nuclease digestion across all human pol II promoters. We identify substantial alterations in nucleosome occupancy over key loci involved in innate immune signaling and other pathways linked to severe disease. Additionally, widespread changes in susceptibility to nuclease digestion are observed in SARS-CoV and SARS-CoV-2 nucleocapsid-expressing cells, particularly over highly expressed genes. These findings suggest that CoV nucleocapsid expression results in remodeled promoter landscapes, potentially contributing to disease outcomes.

We identified 119 to 361 genes with repositioned nucleosomes in their promoter regions, with most repositioning events occurring in a single nucleocapsid-expressing cell line. Gene ontology analysis linked these promoters to innate immune signaling, coagulation, hormone responses, and insulin signaling pathways across multiple CoV samples, with specific disruptions observed at known disease-associated loci such as PABPC1 (76, 109). Additionally, 412 to 2,548 promoters exhibited increased nucleosome occupancy compared to empty vector controls, with gene ontology analysis highlighting enrichment in TGFβ, WNT, hedgehog, and estrogen signaling pathways—all of which are implicated in CoV disease pathology (110–113). Conversely, 134 to 1,717 promoters showed decreased nucleosome occupancy, primarily affecting genes associated with P53 signaling and TNFα pathways, both of which have known roles in CoV disease progression (113–117). These results are especially interesting when we consider that nucleosomes can function as barriers (41) and removal of nucleosomes can lead to aberrant expression of otherwise silenced genes (41, 42). This aligns with previous findings that SARS-CoV-2 nucleocapsid significantly upregulates P53 and TNFα signaling in a reductionist model (119) as we would predict from our nucleosome data.This suggests that nucleosome occupancy changes may precede transcriptional alterations in key disease-relevant pathways.

We also observed that susceptibility to nuclease digestion was moderately impacted by common cold CoV nucleocapsids but was severely disrupted in SARS-CoV and SARS-CoV-2 samples. The most striking alterations were observed in a highly active promoter class characterized by a positioned sensitive region at the transcription start site (TSS), which was absent in SARS-CoV and SARS-CoV-2 samples. Genes in this cluster are associated with ribosomal function and the spliceosome, both of which are directly or indirectly affected by SARS-CoV and SARS-CoV-2 (120–122). In SARS-CoV-2 nucleocapsid-expressing cells, promoters linked to olfactory transduction and viral myocarditis shifted from a silent, diffusely sensitive class to a more active, resistant state, suggesting possible gene de-repression with potential implications for anosmia and myocarditis associated with SARS-CoV-2 infection (66, 123–126). Additionally, promoters transitioning from sensitive or mixed susceptibility patterns to a resistant state in SARS-CoV-2 nucleocapsid-expressing cells were associated with Myc signaling, protein secretion, oxidative phosphorylation, and mTORC1 signaling—all of which are dysregulated in SARS-CoV-2 infections (127–131). These findings indicate that pandemic CoVs induce widespread alterations in promoter susceptibility to nuclease digestion and suggest that these chromatin changes may contribute to disease phenotypes via gene dysregulation.

Our study is the first to evaluate the effects of SARS-CoV-2 and other human CoV nucleocapsids on nucleosome distribution and nuclease susceptibility across all human pol II promoters. We demonstrate that nucleocapsids impact promoter architecture in a reductionist system, with disease-relevant gene classes being particularly affected. Furthermore, common cold CoVs exhibit similar chromatin effects, while SARS-CoV and SARS-CoV-2 nucleocapsids induce more pronounced disruptions in nuclease susceptibility. Beyond providing deeper insights into the functional impact of CoV nucleocapsid proteins on chromatin remodeling, our findings offer a framework for future studies investigating other viral proteins or nucleocapsids from additional viral variants.

## Methods

### Plasmid design and molecular cloning

Mammalian expression vectors encoding nucleocapsid genes from SARS-CoV-2, OC43, NL63, HKU1, and 229E, driven by a CMV promoter, were obtained from the Ginkgo Bioworks COVID-19 Plasmid Collection via Addgene (Table 1). Custom plasmids for MERS-CoV and SARS-CoV nucleocapsids were synthesized by Twist Biosciences using the same backbone. An empty vector control was generated by Q5-directed mutagenesis (New England BioLabs, #E0554S) of the NL63 construct to remove the nucleocapsid gene. All plasmids were validated via Sanger sequencing of the entire nucleocapsid gene.

### Tissue culture and nuclei isolation

A549 lung epithelial cells (ATCC, CCL-185) were cultured in F-12K media (Gibco) supplemented with 10% fetal bovine serum and 1% penicillin-streptomycin and maintained at 37°C with 5% CO_2_. Cells were transfected with nucleocapsid or empty vector plasmids (Table 1) using Turbofect (Thermo Scientific) following the manufacturer’s protocol. Stably transfected clones were selected with puromycin (Gibco), screened by PCR, and confirmed by Sanger sequencing.

For chromatin analysis, cells were crosslinked with 1% v/v formaldehyde (Sigma-Aldrich) and incubated for 10 minutes at room temperature while rocking. Excess formaldehyde was quenched with 125mM glycine and cells were washed twice in PBS. Cells were resuspended in nuclei isolation buffer (10 mM HEPES at pH 7.8, 2 mM MgOAc_2_, 0.3 M sucrose, 1 mM CaCl_2_ and 1% Triton-X) and pelleted by centrifugation at 1000 × *g* for 10 min at 4 °C. Nuclei were washed twice in nuclei isolation buffer and nuclei purity was confirmed by DAPI staining.

### MNase cleavage and preparation of DNA libraries

Each condition was processed in biological replicates. For each reaction, 2.5 × 10^6^ nuclei were resuspended in 500 µL of nuclei isolation buffer and digested with Micrococcal Nuclease (MNase) (Worthington) under light (5U) or heavy (200U) conditions for 10 minutes at 37°C. Digestion was stopped with 50 mM EDTA. Protein-DNA crosslinks were reversed by treating MNase-digested nuclei with 0.2 mg/mL proteinase K and 1% SDS, followed by overnight incubation at 65°C. DNA was purified using a guanidinium-isopropanol and ethanol on a silica column.

Mononucleosomal and sub-mononucleosomal DNA (<200 bp) was isolated from a 2% agarose gel and purified by glycogen precipitation. MNase sequencing libraries were prepared using the Accel-NGS®2S Plus DNA Library Kit (Swift) for Illumina®, with a minimum of 10 ng input DNA per digestion. Libraries were indexed using the Accel-NGS 2S Unique Dual Indexing Kit, normalized with Normalase, and quality-checked via D1000 Tapestation. Library molar concentrations were determined via KAPA qPCR and size-corrected using Tapestation data.

### Solution-based sequence capture and Illumina sequencing

We used a previously validated Roche NimbleGen SeqCap EZ Library (97, 98) to capture 2kb regions flanking the transcription start site (TSS) of all human genes. TSS sequences were repeat masked with human COT DNA, and library adapters were blocked using universal oligos. Biotinylated capture oligos were hybridized to input libraries for 48 hours and purified with streptavidin beads. Enriched sequences were amplified via 15-cycle PCR using TruSeq primers. Final libraries were assessed via Tapestation, and molar concentrations were determined using KAPA qPCR. Paired-end 50 bp sequencing was performed on an Illumina NovaSeq 6000 at the FSU College of Medicine.

### Nucleosome data processing

Raw reads were demultiplexed with Cassava, and adapter sequences and low-quality reads were removed using Trimmomatic (version 0.39) (99). BSequences were aligned to the T2T human genome using Bowtie2 with the --end-to-end --no-mixed --no-discordant flags (100). Mitochondrial and duplicate reads were filtered using Samtools (version 1.10) and Picardtools (version 2.27.3) (101). Read-depth normalization was performed using Picardtools, based on the number of paired-end reads mapping to a 2 kb window centered on the TSS of RefSeq genes. Normalized light and heavy digests were merged using Picardtools to generate total nucleosome profiles. Nucleosome footprints for merged, light, and heavy digests were calculated using DANPOS (version 3.1.1) (102). The log_2_ ratio of light to heavy digests was computed at each time point to determine nucleosome sensitivity using deepTools (version 3.5.1) (103).

Nucleosome distribution changes were extracted from DANPOS output using a custom R script with defined cutoffs for repositioning and occupancy alterations. Repositioned nucleosomes were defined as shifts in summit positions >80 bp between conditions. Changes in nucleosome occupancy were identified as log_2_ fold changes >5 with a p-value and FDR <0.05 compared to empty vector controls. Only nucleosomes consistently altered across replicates were considered high-confidence and used for further analysis. Statistically significant sensitive and resistant nucleosomes were extracted using another custom R script. Sensitivity was defined by a log_2_ fold change in summit score >1.0, while resistance was defined by a fold change <−1.0, with a p-value and FDR <0.05.

### Data Visualization and Functional Analysis

Individual loci visualizations were generated using the PlotGardener package (version 1.4.1) in R (version 4.3.2) and the UCSC Genome Browser (104, 105). Additional processing of bigwig files for upset plots was conducted using R (version 4.3.2), bash, and Python3 with custom scripts. Gene ontology enrichment analyses were performed using Enrichr (106) and visualizations were generated with custom R scripts. Publicly available data multi-omics data for A549 cells was obtained from the Encode portal (107, 108) (https://www.encodeproject.org/) with the following identifiers: ENCFF872SDF, ENCFF985FHV, ENCFF541BTE, ENCFF627QMV, ENCFF369ZNM, ENCFF216KUI, ENCFF556OVF, ENCFF425LVX, ENCFF074PND, and ENCFF774RVE.

### Data availability

All raw and processed nucleosome occupancy and susceptibility to nuclease files were uploaded to the UCSC genome browser and the NCBI GEO database under accession number GSE290042 (https://www.ncbi.nlm.nih.gov/geo/query/acc.cgi?acc=GSE290042). Basic code for computational analysis from file trimming to nucleosome calling has been provided as a supplemental file (Supp. file 1).

## Acknowledgements

We thank Lorea Arambarri, Shahin Behrouz Sharif, and Mahdi Khadem for constructive conversations regarding experimental design and data analysis. This work was supported by the FSU Collaborative Collision COVID-19 Seed Fund. The funders had no role in study design, data collection, data interpretation, or decision to publish this work.

