## Supplementary material for "SARS-CoV-2 nucleocapsid uniquely disrupts chromatin over pathophysiologically relevant gene promoters": Supp_figures

# A

```
Consensus    MSXX-G-X---RXAP-------R-VXWADXSDST-XNQNRGRK-AXPKX------PQ-GXPNXTYSWFXGLT-QHGK-EPLKFPPGQGVPINXXSXP   76
MERS         MASP-------AAP-------AVSFADNNDITNTNLSRGRG-RNPK------PR-AAPNNTVSWYTGLT-QHGK-VPLTFPPGQGVPLNANSTPAQN  74
229E         MAT-------------------VKWADASEPQ-----RGRQ-------------G--RIPYSLYSPLL-VDSE-QPWKVIPRNIVPINKKD-KNKL  54
NL63         MAS-------------------VNWADDRA--------ARK-K--------------FPPPSFYMPLLVSSDK-APYRVIPRNIVPIGKGN-KDEQ  52
HKU1         MSYTPGHYAGSRSSSGNRSGILKKTSWADQSERNYQTFNRGRK-TQPKFTVS-TQPQ-GNTIPHYSWFSGIT-QFQKGRDFKFSDGQGVPIAFGVPPSEA  96
OC43         MSFTPGKQSSSRASSGNRSGN-GILKWADQSDQFRNVQTRGRR-AQPKQTATSQQPSGGNVVPYYSWFSGIT-QFQKGKEFEFVEGQGVPIAPGVPATEA  97
SARS-CoV-2   MSDN-GPQN-QRNAP-------R-ITFGGPSDSTGSNQNGERSGARSKQ----RRPQ-GLPNNTASWFTALT-QHGK-EDLKFPRGQGVPINTNSSPDDQ  83
SARS-CoV     MSDN-GPQSNQRSAP-------R-ITFGGPTDSTDNNQNGGRNGARPKQ----RRPQ-GLPNNTASWFTALT-QHGK-EELRFPRGQGVPINTNSGPDDQ  84

Consensus    IGYWRRQTR-XFRTGDGKXKXLSPRWYFYYLGTGPXADLPYGAXKDGVVWVATEGAX-NTPXD-IGTRNPNNBXAIXTQFPPGTXLPKGFY-VEGS-    171
MERS         AGYWRRQDR-KINTGNG-IKQLAPRWYFYYTGTGPEAALPFRAVKDGIVWVHEDGAT-DAPST-FGTRNPNNDSAIVTQFAPGTKLPKNFH-IEGTGGNS  169
229E         IGYWNVQKR--FRTRKGKRVDLSPKLHFYYLGTGPHKDAKFRERVEGVVWVAVDGAK-TEPTG-YGVRRKNSEPEIP-HFN--QKLPNGVT-VVEE-PDS  145
NL63         IGYWNVQER--WRMRRGQRVDLPPKVHFYYLGTGPHKDLKFRQRSDGVVWVAKEGAK-TVNTS-LGNRKRNQKP-LEPKFS--IALPPELSVVEFE-DRS  144
HKU1         KGYWYRHSRRSFKTADGQQKQLLPRWYFYYLGTGPYANASYGESLEGVFWVANHQADTSTPSD-VSSRDPTTQEAIPTRFPPGTILPQGYY-VEGS-GRS  193
OC43         KGYWYRHNRRSFKTADGNQRQLLPRWYFYYLGTGPHAKDQYGTDIDGVYWVASNQADVNTPAD-IVDRDPSSDEAIPTRFPPGTVLPQGYY-IEGS-GRS  194
SARS-CoV-2   IGYYRRATR-RIRGGDGKMKDLSPRWYFYYLGTGPEAGLPYGANKDGIIWVATEGAL-NTPKDHIGTRNPANNAAIVLQLPQGTTLPKGFY-AEGSRGGS  180
SARS-CoV     IGYYRRATR-RVRGGDGKMKELSPRWYFYYLGTGPEASLPYGANKEGIVWVATEGAL-NTPKDHIGTRNPNNNAATVLQLPQGTTLPKGFY-AEGSRGGS  181

Consensus    QASSRSSSRS--RNSSRNSSXGS-SRGNSPXRMASSGX------AVALALL--LLDXLNQLXSX-VSGK-GKQ-XXP-----------------QX   239
MERS         QSSSRASSVS--RNSSRSSSQGS-RSGNS-TRGTSPGP--SGIGAVGGDLL--YLDLLNRLQAL-ESGK-VKQ-SQP-----------------KVITK  240
229E         RAPSRSQSRS--QSRGRGESKPQ-SRNPSSDRNHNSQD--DIMKAVAAALKSLGFDKPQEKDKK-SAKT-GTP-KPSRNQSPASSQTSAKSLARSQSSET  237
NL63         NNSSRASSRSSTRNNSRDSSRST-SRQQSRTRSDSNQSSSDLVAAVTLALK--NLGFDNQSKSPSSSGT-STP-KKP--NKP--------------LSQPR  224
HKU1         ASNSRPGSRS--QSRGPNN-RSL-SRSNSNFRHSDSIV-------------KPDMADEIANL-VLAKLGKD-SKP-----------------QQVTK  254
OC43         APNSRSTSRT----SSRASSAGSRSRANSGNRTPTSGV-------------TPDMADQIASL-VLAKLGKDATKP-----------------QQVTK  256
SARS-CoV-2   QASSRSSSRS--RNSSRNTPGS-SRGTSPARMAGNGG------DAALALL--LLDRLNQLESK-MSGK-GQQ-QQG-----------------QTVTK  248
SARS-CoV     QASSRSSSRS--IGNSRNSTPGS-SRGNSPARMASGGG------ETALALL--LLDRLNQLESK-VSGK-QQ-QQG-----------------QTVTK  249

Consensus    KXAXEA----LKKPRQKRTPTKQ--XNVTQCFGXRGP---QGNFGDXELXKLGTDDKXXPQJAELAPXASAFFFMSRIGLEXXX----------DV   319
MERS         KDAAAA----KNKMRHKRTSTKS--FNMVQAFGLRGPGDLQGNFGDLQLTKLGTEDPRWPQIAELAPTASAFMGMSQFKLTHQNN-----DDHGNPVYFL  329
229E         KEQKHE----MQKPRWKRQPNDDVTSNVTQCFGPRDL---DHNFGSAGVVANGVKAKGYPQFAELVPSTAAMLFDSHIVSKESG-----------NTVVL  319
NL63         ADKPSQ----LKKPRWKRVPTRE--ENVIQCFGPRDF---NHNMGDSDLVQNGVDAKGFPQLAELIPNQAALFFDSEVSTDEVG-----------DNVQI  304
HKU1         QNAKEIRHKILTKPRQKRTPNKH--CNVQQCFGKRGP---SQNFGNAEMLKLGTNDPQFPILAELAPTPGAFFFGSKLDLVKRDS---EADSPVKDVFEL  346
OC43         HTAKEVRQKILNKPRQKRSPNKQ--CTVQQCFGKRGP---NQNFGGGEMLKLGTSDPQFPILAELAPTAGAFFFGSRLELAKVQNLSGNPDEPQKDVYEL  351
SARS-CoV-2   KSAAEA----SKKPRQKRTATKA--YNVTQAFGRRGPEQTQGNFGDQELIRQGTDYKHWPQIAQFAPSASAFFGMSRIGMEVTP-----------SGTWL  331
SARS-CoV     KSAAEA----SKKPRQKRTATKQ--YNVTQAFGRRGPEQTQGNFGDQDLIRQGTDYKHWPQIAQFAPSASAFFGMSRIGMEVTP-----------SGTWL  332

Consensus    TYTGAIKLDXKXPNFXKXXXELLNXNIDAYKTFP--XPKXX--KXKXTDXXSXXPQRQR--------------------Q-XXTLLPAA--X--XX  388
MERS         RYSGAIKLDPKNPNYNKWLELLEQNIDAYKTFPKKEKKQKAPKEESTDQMSEPPKEQR--------------------VQGSITQR-------TRTRP  400
229E         TFTTRVTVPKDHPHLGKFLEELNAFTREMQQHPLLNPSALEFNPSQTSPATAEPVRDE-------------------------------V------  378
NL63         TYTYKMLVAKDNKNLPKFIE----QISAFTK-----PSSI--KEMQSQS-SHVAQN-----------TVLNASIPE-------SKPLADDD------  365
HKU1         HYSGSIRFDSTLPGFETIMKVLEENLNAYVNSN------------QNTDSDSLSSKPQRKRGVKQLPEQFDSLNLSAGT----QHISNDFTPE-----DHSLL  428
OC43         RYNGAIRFDSTLSGFETIMKVLNENLNAYQQQ-------------DGMMNMSPKPQRQRGHKNGQGENDNISVAVPKSRVQQNKSRELTAE-----DISLL  434
SARS-CoV-2   TYTGAIKLDDKDPNFKDQVILLNKHIDAYKTFPPTEPKKD--KKKKADETQALPQRQK--------------------KQQTVTLLPAADLDDFSKQLQ  408
SARS-CoV     TYHGAIKLDDKDPQFKDNVILLNKHIDAYKTFPPTEPKKD--KKKKTDEAQPLPQRQK--------------------KQPTVTLLPAADMDDFSRQLQ  409

Consensus    -SMDGPXX-DXXXA      400
MERS         SVQPGPMI-DVNTD      413
229E         -SIETDII--DEVN      389
NL63         -SAIIEIV-NEVLH      377
HKU1         ATLDDPYV-EDSVA      441
OC43         KKMDEPYTEDTSEI      448
SARS-CoV-2   QS--MSSA-DSTQA      419
SARS-CoV     NSMSGASA-DSTQA      422
```

### Supp. Fig. 1. Alignment of CoV nucleocapsid proteins

A. Clustal Omega alignment of CoV nucleocapsid proteins. Residues which differ from the consensus are highlighted in grey.

A

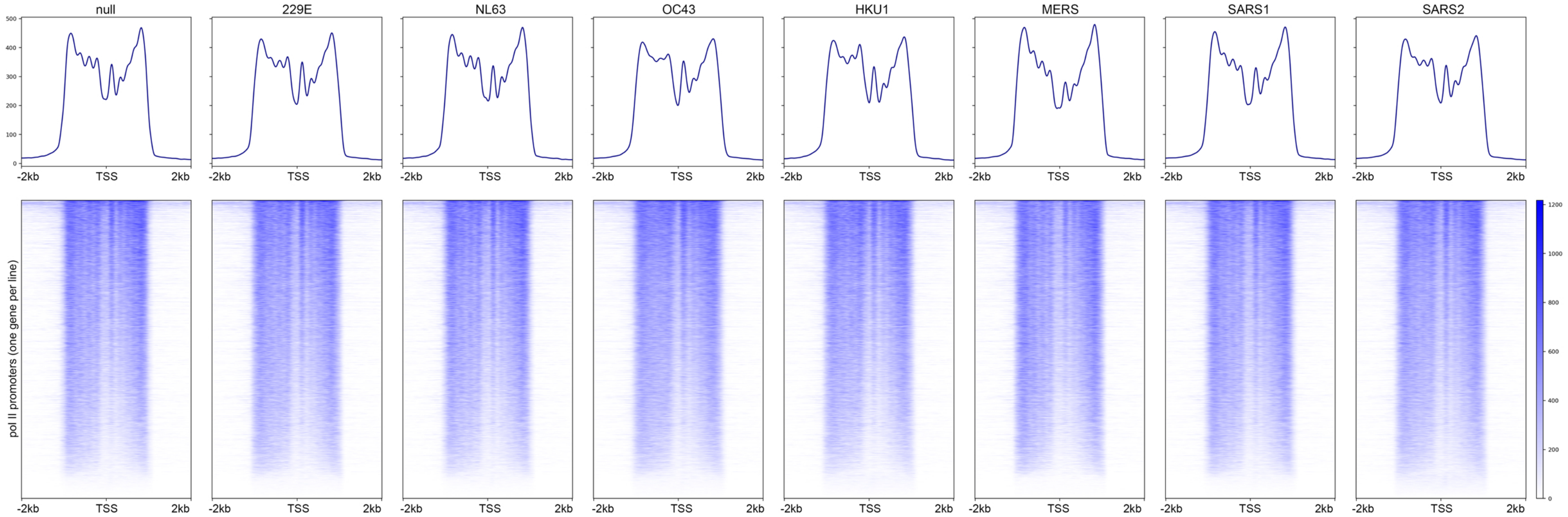
null
229E
NL63
OC43
HKU1
MERS
SARS1
SARS2
-2kb
TSS
2kb
pol II promoters (one gene per line)

### Supp. Fig. 2. Sequence capture effectively enriches human promoter regions

A. Heatmaps of nucleosome distribution (blue) over a 4kb region centered on the TSS of pol II promoters (one gene per line) with a line plot showing average signal above each plot. The overall gene order is identical across heatmaps and is determined by maximum signal values.

**A**

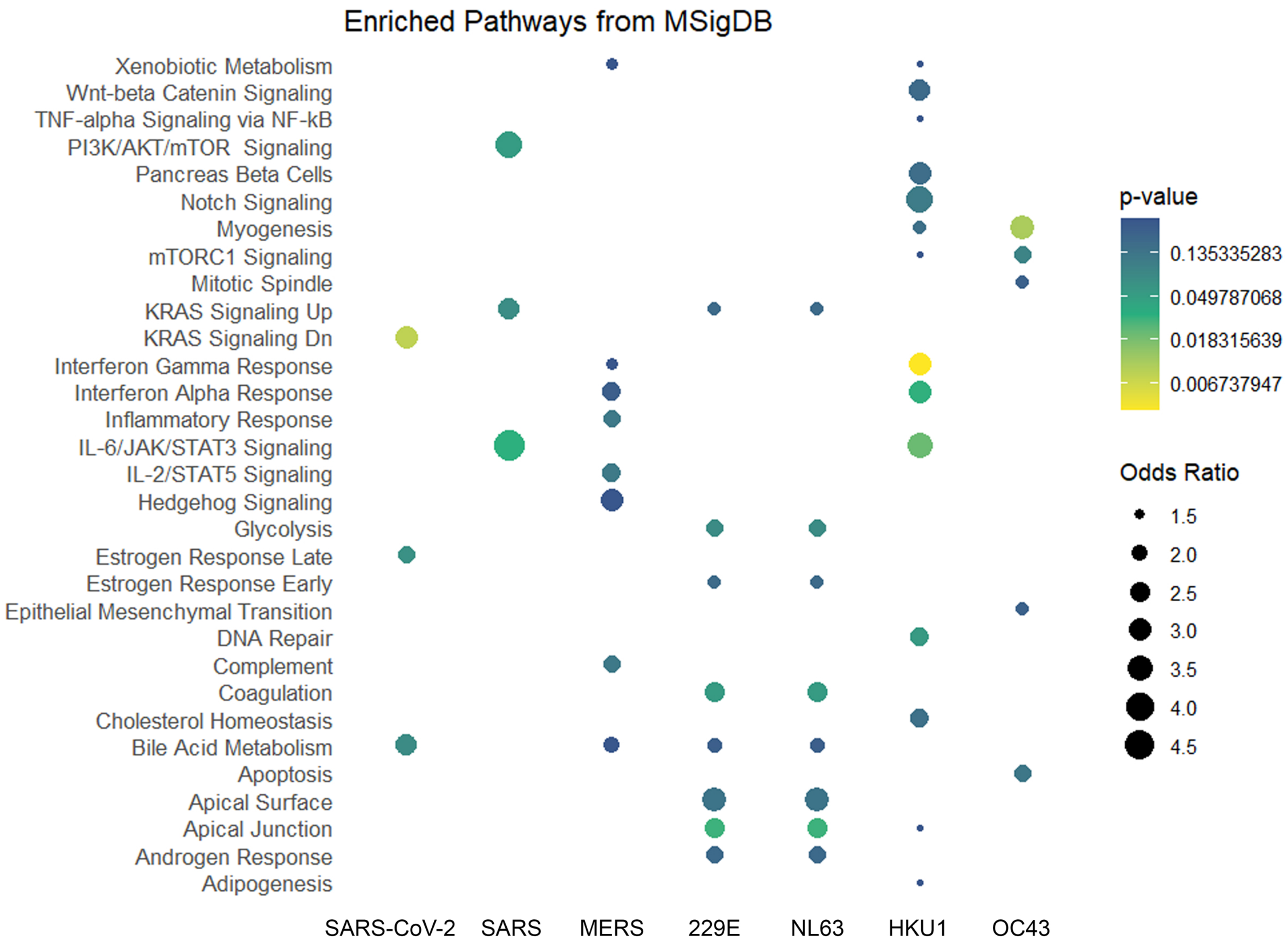
Enriched Pathways from MSigDB
Xenobiotic Metabolism
Wnt-beta Catenin Signaling
TNF-alpha Signaling via NF-kB
PI3K/AKT/mTOR Signaling
Pancreas Beta Cells
Notch Signaling
Myogenesis
mTORC1 Signaling
Mitotic Spindle
KRAS Signaling Up
KRAS Signaling Dn
Interferon Gamma Response
Interferon Alpha Response
Inflammatory Response
IL-6/JAK/STAT3 Signaling
IL-2/STAT5 Signaling
Hedgehog Signaling
Glycolysis
Estrogen Response Late
Estrogen Response Early
Epithelial Mesenchymal Transition
DNA Repair
Complement
Coagulation
Cholesterol Homeostasis
Bile Acid Metabolism
Apoptosis
Apical Surface
Apical Junction
Androgen Response
Adipogenesis
SARS-CoV-2
SARS
MERS
229E
NL63
HKU1
OC43
p-value
0.135335283
0.049787068
0.018315639
0.006737947
Odds Ratio
1.5
2.0
2.5
3.0
3.5
4.0
4.5

### Supp. Fig. 3. Repositioned nucleosomes in CoV nucleocapsid expressing cell lines are associated with inflammatory and hormone responses

A. Dot plot of top MSigDB Hallmark Gene Ontology terms for genes with one or more repositioned nucleosomes in each CoV sample.

A

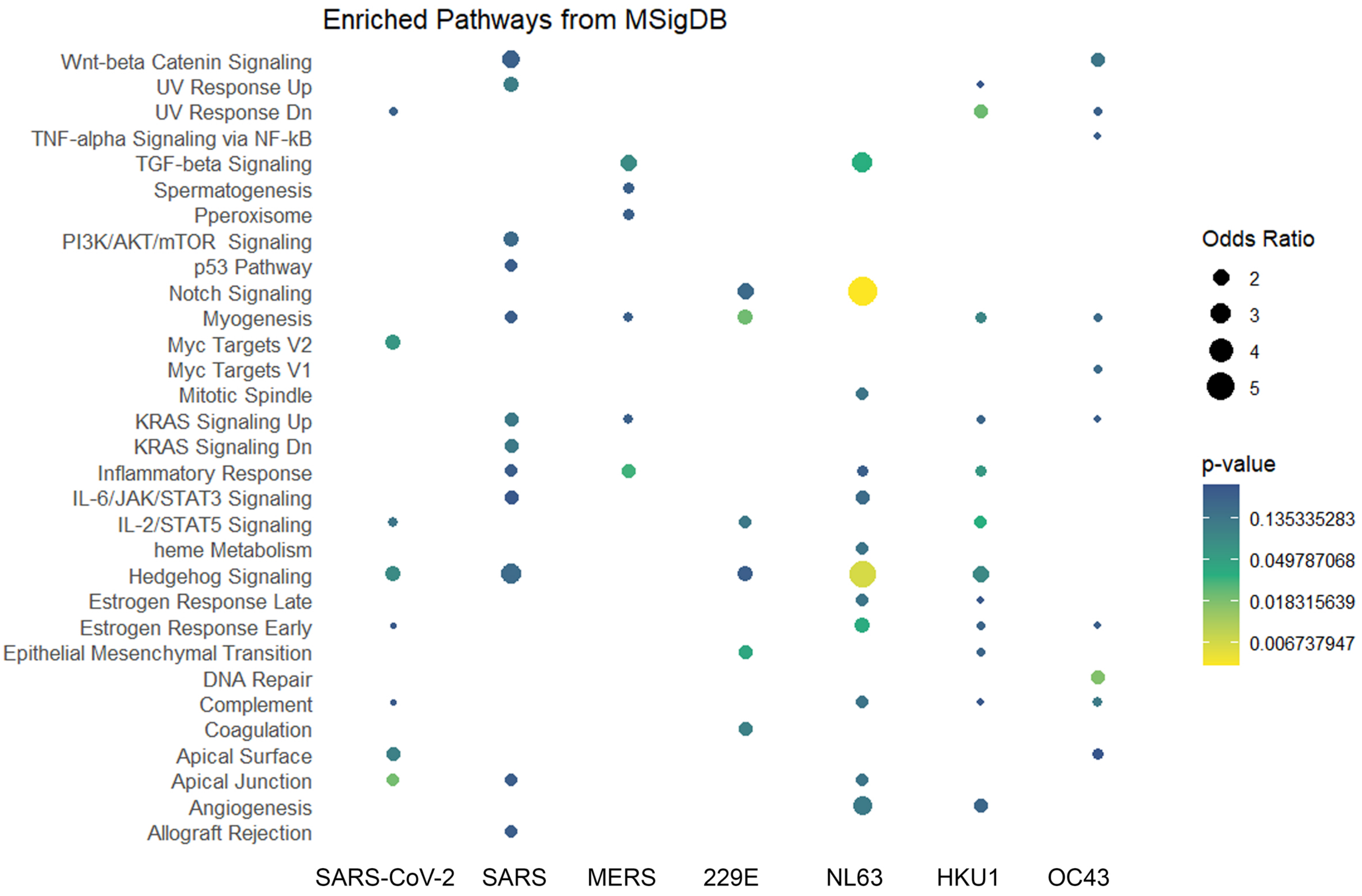
Enriched Pathways from MSigDB
Wnt-beta Catenin Signaling
UV Response Up
UV Response Dn
TNF-alpha Signaling via NF-kB
TGF-beta Signaling
Spermatogenesis
Pperoxisome
PI3K/AKT/mTOR Signaling
p53 Pathway
Notch Signaling
Myogenesis
Myc Targets V2
Myc Targets V1
Mitotic Spindle
KRAS Signaling Up
KRAS Signaling Dn
Inflammatory Response
IL-6/JAK/STAT3 Signaling
IL-2/STAT5 Signaling
heme Metabolism
Hedgehog Signaling
Estrogen Response Late
Estrogen Response Early
Epithelial Mesenchymal Transition
DNA Repair
Complement
Coagulation
Apical Surface
Apical Junction
Angiogenesis
Allograft Rejection
SARS-CoV-2
SARS
MERS
229E
NL63
HKU1
OC43
Odds Ratio
2
3
4
5
p-value
0.135335283
0.049787068
0.018315639
0.006737947

### Supp. Fig. 4. Nucleosomes with increased occupancy in CoV nucleocapsid expressing cell lines are associated with signaling pathways and inflammatory responses

A. Dot plot of top MSigDB Hallmark Gene Ontology terms for genes with one or more nucleosomes with increased occupancy in each CoV sample.

A

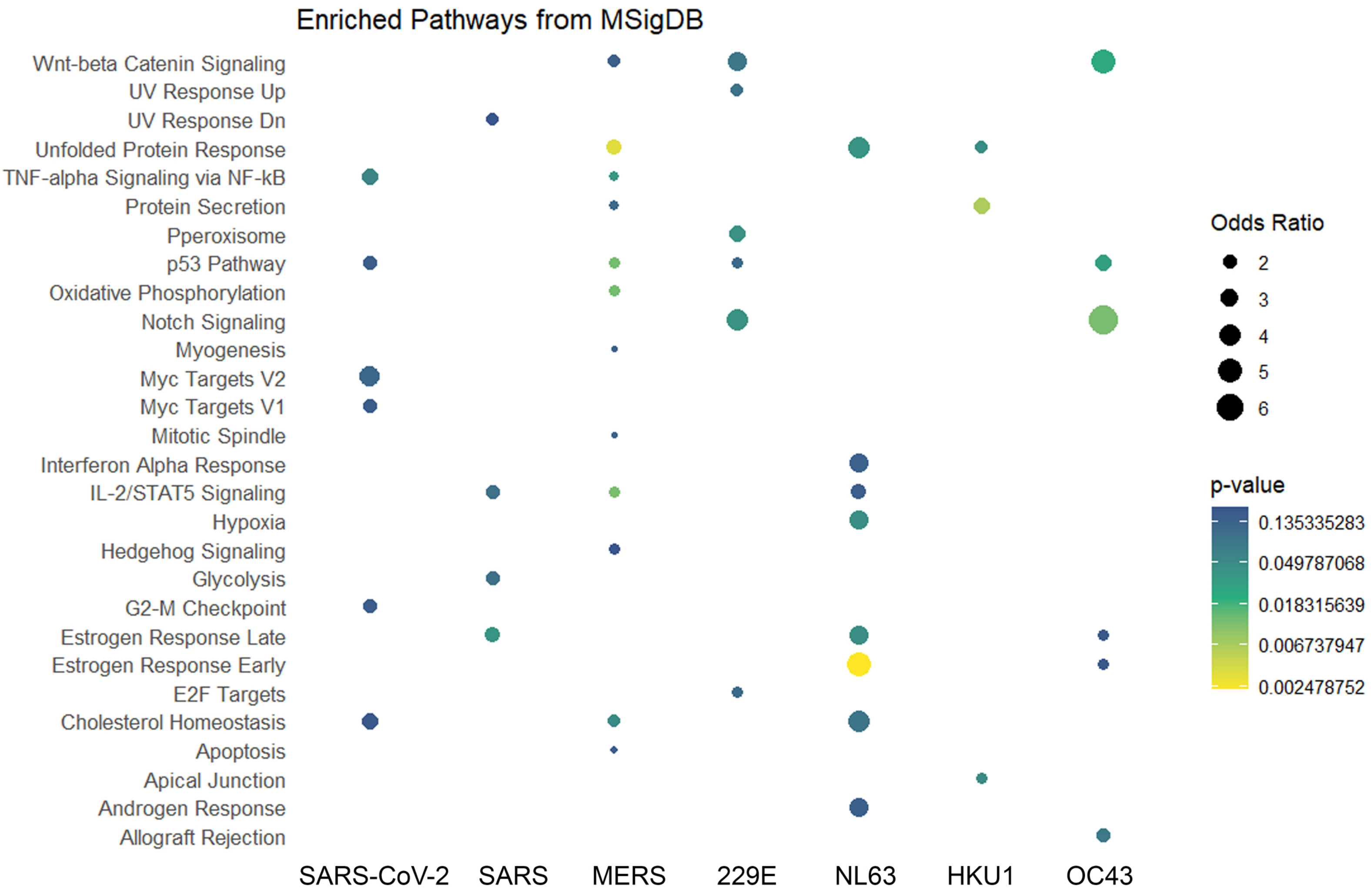
Enriched Pathways from MSigDB
Wnt-beta Catenin Signaling
UV Response Up
UV Response Dn
Unfolded Protein Response
TNF-alpha Signaling via NF-kB
Protein Secretion
Pperoxisome
p53 Pathway
Oxidative Phosphorylation
Notch Signaling
Myogenesis
Myc Targets V2
Myc Targets V1
Mitotic Spindle
Interferon Alpha Response
IL-2/STAT5 Signaling
Hypoxia
Hedgehog Signaling
Glycolysis
G2-M Checkpoint
Estrogen Response Late
Estrogen Response Early
E2F Targets
Cholesterol Homeostasis
Apoptosis
Apical Junction
Androgen Response
Allograft Rejection
SARS-CoV-2
SARS
MERS
229E
NL63
HKU1
OC43
Odds Ratio
2
3
4
5
6
p-value
0.135335283
0.049787068
0.018315639
0.006737947
0.002478752

### Supp. Fig. 5. Nucleosomes with decreased occupancy in CoV nucleocapsid expressing cell lines are associated with signaling pathways and hormone responses

A. Dot plot of top MSigDB Hallmark Gene Ontology terms for genes with one or more nucleosomes with decreased occupancy in each CoV sample.

A

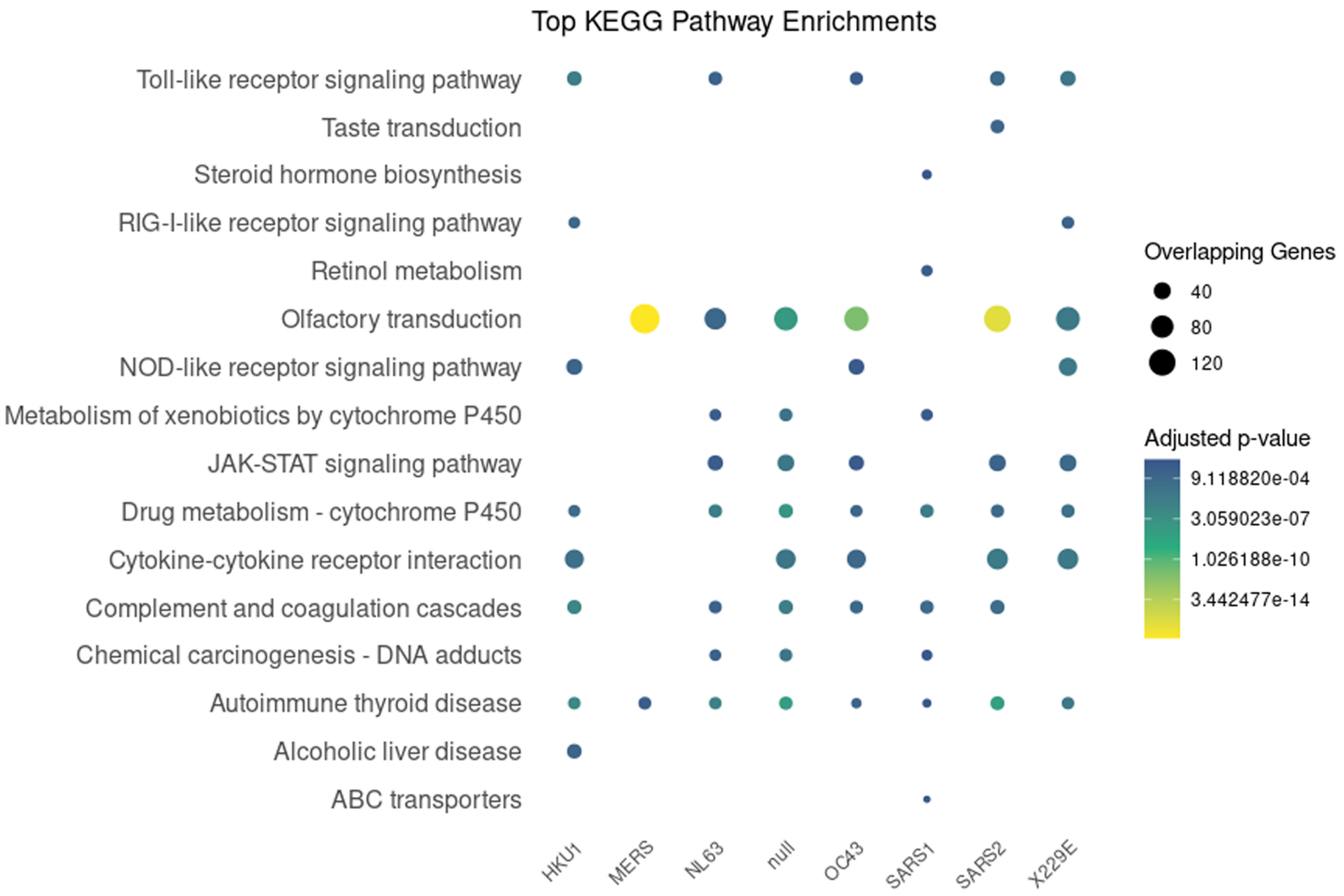

Top KEGG Pathway Enrichments
Toll-like receptor signaling pathway
Taste transduction
Steroid hormone biosynthesis
RIG-I-like receptor signaling pathway
Retinol metabolism
Olfactory transduction
NOD-like receptor signaling pathway
Metabolism of xenobiotics by cytochrome P450
JAK-STAT signaling pathway
Drug metabolism - cytochrome P450
Cytokine-cytokine receptor interaction
Complement and coagulation cascades
Chemical carcinogenesis - DNA adducts
Autoimmune thyroid disease
Alcoholic liver disease
ABC transporters
HKU1
MERS
NL63
null
OC43
SARS1
SARS2
X229E
Overlapping Genes
40
80
120
Adjusted p-value
9.118820e-04
3.059023e-07
1.026188e-10
3.442477e-14


B

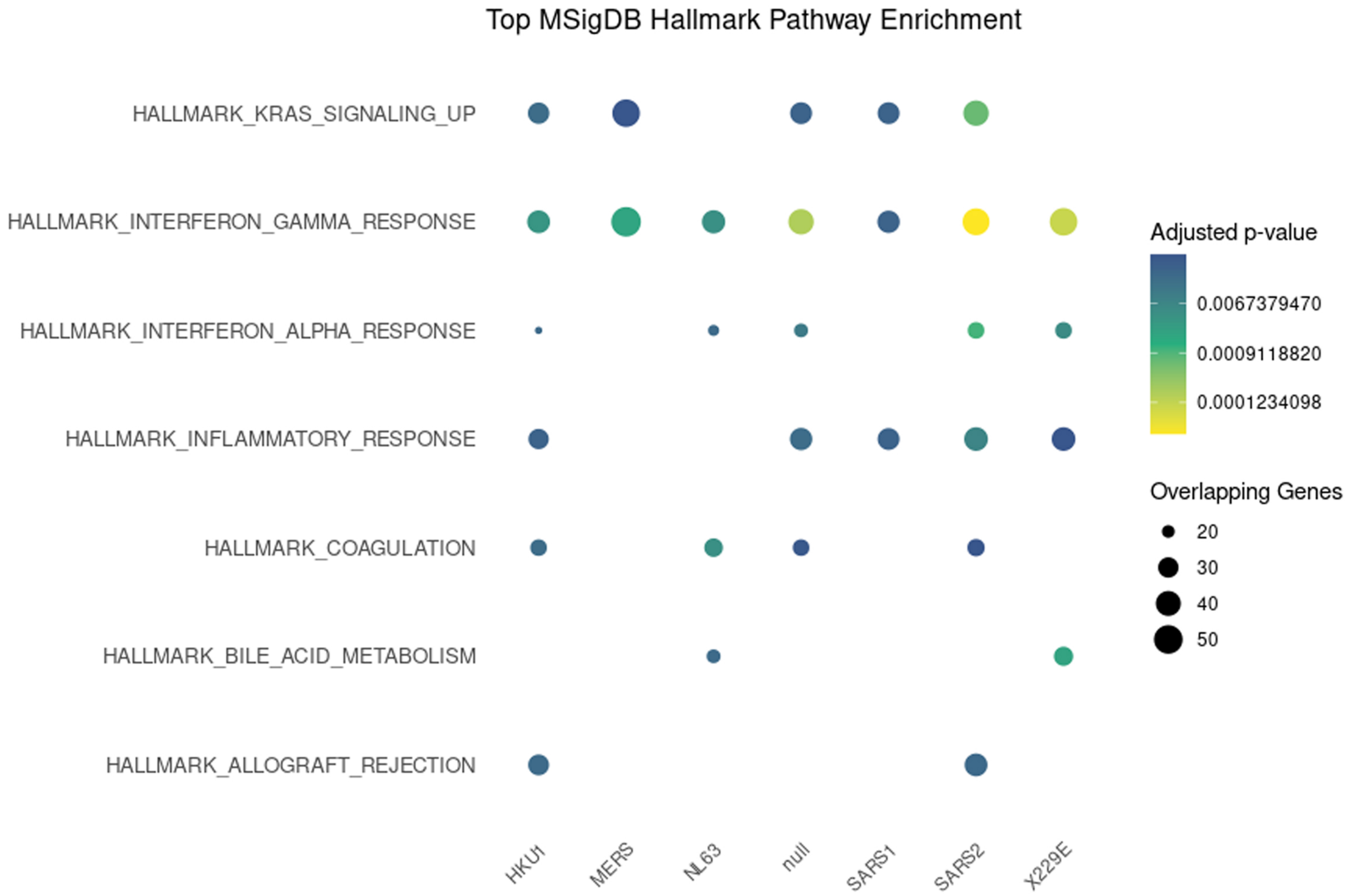

Top MSigDB Hallmark Pathway Enrichment
HALLMARK_KRAS_SIGNALING_UP
HALLMARK_INTERFERON_GAMMA_RESPONSE
HALLMARK_INTERFERON_ALPHA_RESPONSE
HALLMARK_INFLAMMATORY_RESPONSE
HALLMARK_COAGULATION
HALLMARK_BILE_ACID_METABOLISM
HALLMARK_ALLOGRAFT_REJECTION
HKU1
MERS
NL63
null
SARS1
SARS2
X229E
Adjusted p-value
0.0067379470
0.0009118820
0.0001234098
Overlapping Genes
20
30
40
50

### Supp. Fig. 6. Nucleosomes within diffusely sensitive clusters in CoV nucleocapsid expressing cell lines are associated with immune signaling and coagulation cascades

A. Dot plot of top KEGG Gene Ontology terms for genes belonging to diffusely sensitive promoter clusters (defined in Fig. 4B). B. Dot plot of top MSigDB Hallmark Gene Ontology terms for genes belonging to diffusely sensitive promoter clusters (defined in Fig. 4B)

A

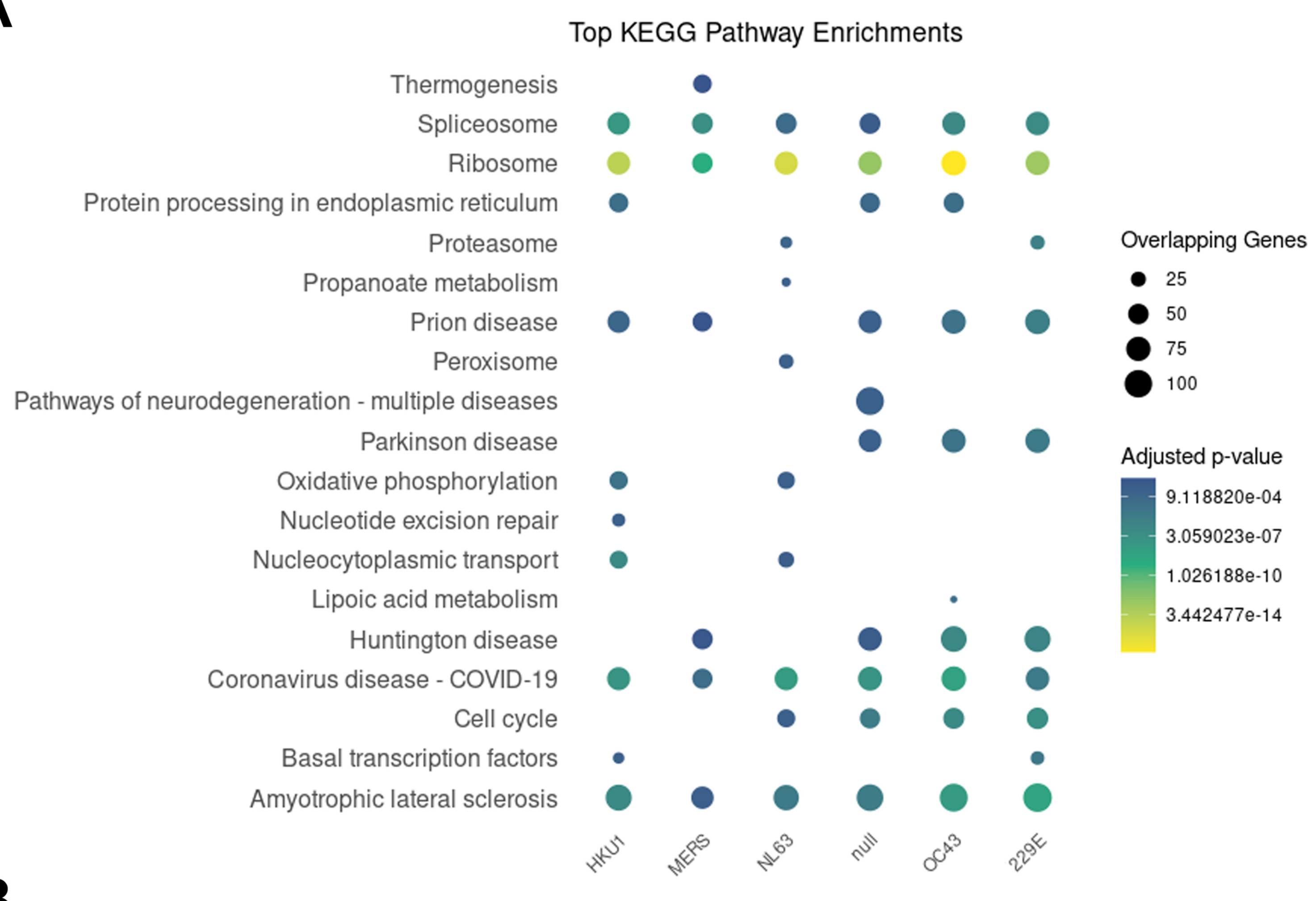

Top KEGG Pathway Enrichments
Thermogenesis
Spliceosome
Ribosome
Protein processing in endoplasmic reticulum
Proteasome
Propanoate metabolism
Prion disease
Peroxisome
Pathways of neurodegeneration - multiple diseases
Parkinson disease
Oxidative phosphorylation
Nucleotide excision repair
Nucleocytoplasmic transport
Lipoic acid metabolism
Huntington disease
Coronavirus disease - COVID-19
Cell cycle
Basal transcription factors
Amyotrophic lateral sclerosis
HKU1
MERS
NL63
null
OC43
229E
Overlapping Genes
25
50
75
100
Adjusted p-value
9.118820e-04
3.059023e-07
1.026188e-10
3.442477e-14


B

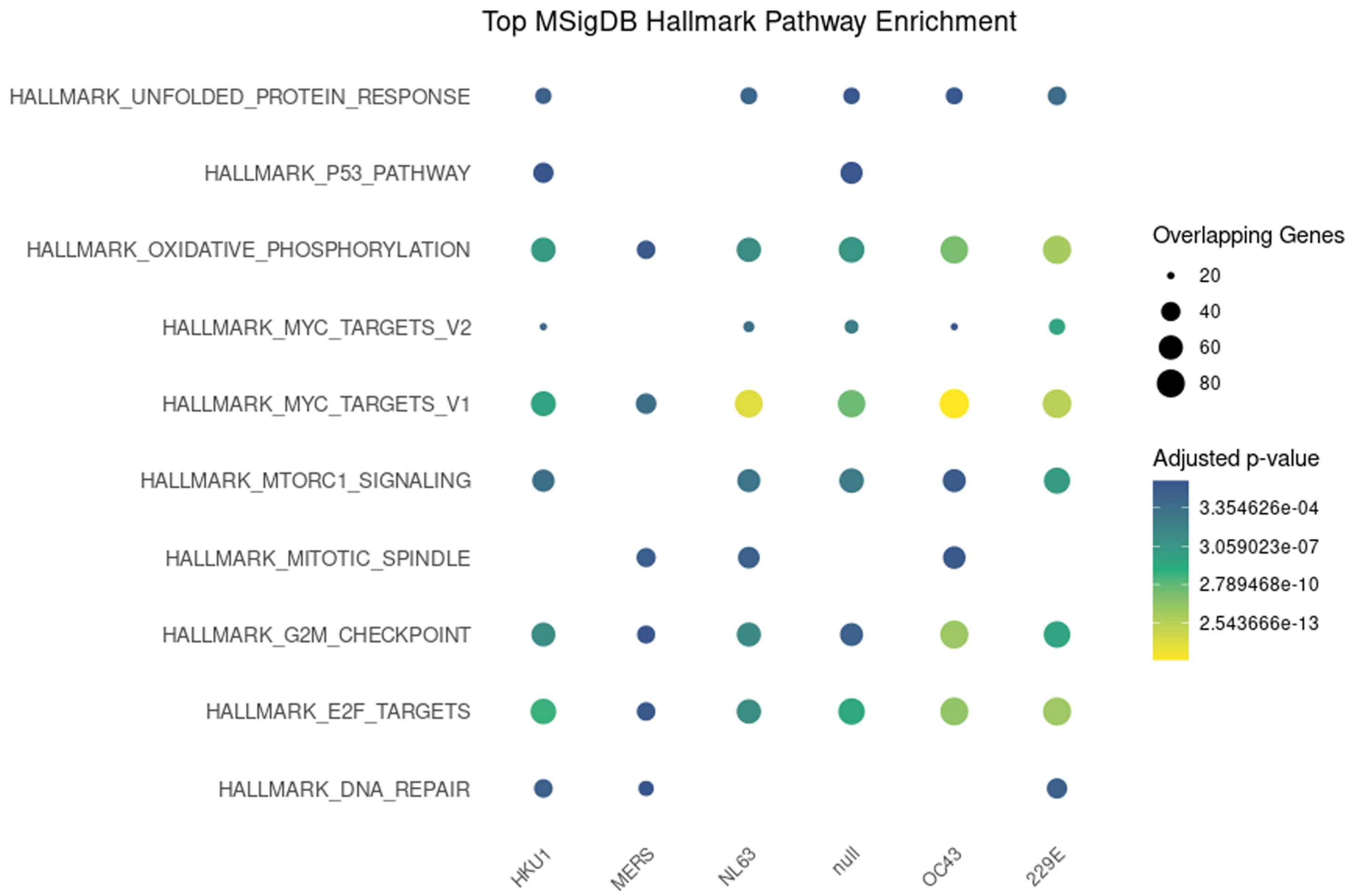

Top MSigDB Hallmark Pathway Enrichment
HALLMARK_UNFOLDED_PROTEIN_RESPONSE
HALLMARK_P53_PATHWAY
HALLMARK_OXIDATIVE_PHOSPHORYLATION
HALLMARK_MYC_TARGETS_V2
HALLMARK_MYC_TARGETS_V1
HALLMARK_MTORC1_SIGNALING
HALLMARK_MITOTIC_SPINDLE
HALLMARK_G2M_CHECKPOINT
HALLMARK_E2F_TARGETS
HALLMARK_DNA_REPAIR
HKU1
MERS
NL63
null
OC43
229E
Overlapping Genes
20
40
60
80
Adjusted p-value
3.354626e-04
3.059023e-07
2.789468e-10
2.543666e-13

### Supp. Fig. 7. Nucleosomes within positioned sensitive clusters over the TSS in CoV nucleocapsid expressing cell lines are associated with RNA splicing and ribosome function

A. Dot plot of top KEGG Gene Ontology terms for genes belonging to positioned sensitive promoter clusters (defined in Fig. 4B). B. Dot plot of top  MSigDB Hallmark Gene Ontology terms for genes belonging to positioned sensitive promoter clusters (defined in Fig. 4B)

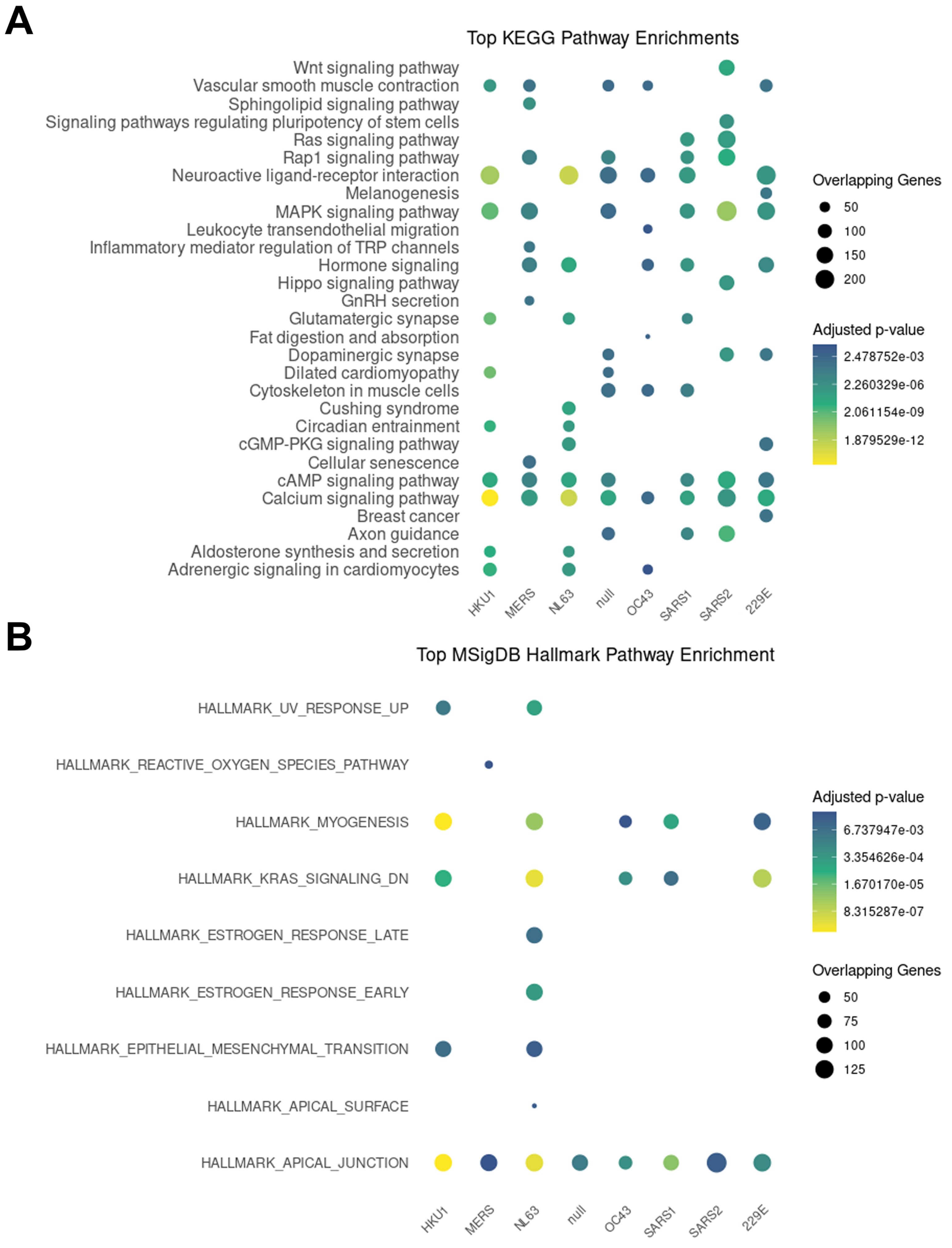
A
Top KEGG Pathway Enrichments
Wnt signaling pathway
Vascular smooth muscle contraction
Sphingolipid signaling pathway
Signaling pathways regulating pluripotency of stem cells
Ras signaling pathway
Rap1 signaling pathway
Neuroactive ligand-receptor interaction
Melanogenesis
MAPK signaling pathway
Leukocyte transendothelial migration
Inflammatory mediator regulation of TRP channels
Hormone signaling
Hippo signaling pathway
GnRH secretion
Glutamatergic synapse
Fat digestion and absorption
Dopaminergic synapse
Dilated cardiomyopathy
Cytoskeleton in muscle cells
Cushing syndrome
Circadian entrainment
cGMP-PKG signaling pathway
Cellular senescence
cAMP signaling pathway
Calcium signaling pathway
Breast cancer
Axon guidance
Aldosterone synthesis and secretion
Adrenergic signaling in cardiomyocytes
HKU1
MERS
NL63
null
OC43
SARS1
SARS2
229E
Overlapping Genes
50
100
150
200
Adjusted p-value
2.478752e-03
2.260329e-06
2.061154e-09
1.879529e-12
B
Top MSigDB Hallmark Pathway Enrichment
HALLMARK_UV_RESPONSE_UP
HALLMARK_REACTIVE_OXYGEN_SPECIES_PATHWAY
HALLMARK_MYOGENESIS
HALLMARK_KRAS_SIGNALING_DN
HALLMARK_ESTROGEN_RESPONSE_LATE
HALLMARK_ESTROGEN_RESPONSE_EARLY
HALLMARK_EPITHELIAL_MESENCHYMAL_TRANSITION
HALLMARK_APICAL_SURFACE
HALLMARK_APICAL_JUNCTION
HKU1
MERS
NL63
null
OC43
SARS1
SARS2
229E
Adjusted p-value
6.737947e-03
3.354626e-04
1.670170e-05
8.315287e-07
Overlapping Genes
50
75
100
125

### Supp. Fig. 8. Nucleosomes within resistant clusters in CoV nucleocapsid expressing cell lines are associated with signaling cascades and hormone responses

A. Dot plot of top KEGG Gene Ontology terms for genes belonging to resistant promoter clusters (defined in Fig. 4B). B. Dot plot of top  MSigDB Hallmark Gene Ontology terms for genes belonging to resistant promoter clusters (defined in Fig. 4B)
